# Regulation of Liprin-α phase separation by CASK is disrupted by a mutation in its CaM kinase domain

**DOI:** 10.1101/2022.04.21.489014

**Authors:** Debora Tibbe, Pia Ferle, Christoph Krisp, Sheela Nampoothiri, Ghayda Mirzaa, Melissa Assaf, Sumit Parikh, Kerstin Kutsche, Hans-Jürgen Kreienkamp

## Abstract

CASK is a unique membrane associated guanylate kinase (MAGUK), due to its Ca^2+^/calmodulin-dependent kinase (CaMK) domain. We describe four male patients with a severe neurodevelopmental disorder with microcephaly carrying missense variants affecting the CaMK domain. One boy who carried the p.E115K variant and died at an early age showed pontocerebellar hypoplasia (PCH) in addition to microcephaly, thus exhibiting the classical MICPCH phenotype observed in individuals with *CASK* loss-of-function variants. All four variants selectively weaken the interaction of CASK with Liprin-α2, a component of the presynaptic active zone. Liprin-α proteins form spherical condensates in a process termed liquid-liquid phase separation (LLPS), which we observe here in Liprin-α2 overexpressing HEK293T cells and primary cultured neurons. Condensate formation is reversed by interaction of Liprin-α2 with CASK; this is associated with altered phosphorylation of Liprin-α2. The p.E115K variant fails to interfere with condensate formation. As the individual carrying this variant had the severe MICPCH disorder, we suggest that regulation of Liprin-α2-mediated LLPS is a new functional feature of CASK which must be maintained to prevent PCH.

## Introduction

Perturbations in synapse formation, synaptic protein complexes and synaptic transmission are associated with neurodevelopmental disorders in humans (Bourgeron, 2015; Grabrucker *et al*, 2011). Both pre- and postsynaptically, large protein complexes are formed which contribute to synaptic architecture and function. One such complex, the presynaptic active zone, consists of a dense network of core constituents ELKS, Liprin-α, RIM, RIM-BP and Munc13. The active zone complex is held together through multiple interactions via highly conserved domains such as C_2_, PDZ and SH3 domains (Sudhof, 2012). Recently, it became clear that active zone assembly also relies on a process termed liquid-liquid phase separation (LLPS), which involves the recruitment of proteins into condensates mediated by multiple intrinsically disordered regions (IDRs) (Emperador-Melero *et al*, 2021; Liang *et al*, 2021; Xie *et al*, 2021).

The active zone is required for the recruitment of synaptic vesicles to release sites precisely opposite to postsynaptic specializations containing the appropriate neurotransmitter receptors (Sudhof, 2012). This positioning requires transsynaptic adhesion complexes such as the Neurexin/Neuroligin pair of adhesion molecules (Sudhof, 2008). Neurexins are linked to the active zone through the Ca^2+^/calmodulin-dependent serine protein kinase CASK (Hata *et al*, 1996). CASK binds to Neurexins via its C-terminal PDZ-SH3-GK (PSG) module (Hata *et al*., 1996; Li *et al*, 2014; Pan *et al*, 2021), whereas the N-terminal CaM-dependent kinase domain (CaMK domain) is involved in protein interactions with Liprin-α (LaConte *et al*, 2016; Wei *et al*, 2011). Additional interactions with Mint1, also through the CaMK domain, and Lin/Veli proteins through the L27.2 domain, establish CASK as a multivalent scaffold protein (Butz *et al*, 1998; Tabuchi *et al*, 2002). The CaMK domain exhibits a Mg^2+^-sensitive, atypical kinase activity which may phosphorylate the Neurexin C-terminus (Mukherjee *et al*, 2008). The functional relevance of this activity is unclear. In addition to its presynaptic role, CASK has been reported to act as a transcriptional regulator during neuronal development (Hsueh *et al*, 2000) and as a regulator of postsynaptic glutamate receptor trafficking (Jeyifous *et al*, 2009).

Loss-of-function variants in the X-chromosomal *CASK* gene lead to microcephaly with pontine and cerebellar hypoplasia (MICPCH) and intellectual disability (ID) in females in the heterozygous state and in males in the hemizygous state. Furthermore, several missense variants have been described which are associated with neurodevelopmental disorders of variable severity. These variants mostly affect males and are often inherited from healthy mothers (Hackett *et al*, 2010; Najm *et al*, 2008; Pan *et al*., 2021).

So far, the pathogenic mechanisms of *CASK* mutations remain unclear; in particular, it is unknown why some variants cause ID, while others are associated also with pontocerebellar hypoplasia (PCH). As CASK fulfils multiple functions at the pre- and postsynapse and in the nucleus, we do not know which of these apparently separate functions contributes most strongly to the patient’s phenotype. We have begun to address this by analysing a larger number of missense variants with respect to interactions with a panel of known CASK associated proteins. Our initial data indicated that the presynaptic role of CASK was affected in most cases, as most variants interfered with Neurexin binding (Pan *et al*., 2021). Here, we identify four male patients carrying missense variants in the CaMK domain of CASK. All four variants selectively interfere with Liprin-α2 binding, strongly supporting a disturbed presynaptic function of CASK as a major pathogenic mechanism. Importantly, we also observe that in human cells and in neurons, Liprin-α2 undergoes formation of spherical condensates. This process can be reversed by interaction with CASK, but not by a CASK variant which is deficient in Liprin-α2 binding. Our data uncover a new aspect of the molecular function of CASK which may be relevant for proper formation of the presynaptic active zone.

## Results

### Missense variants altering the CaMK domain of CASK

Four novel hemizygous missense variants (p.E115K, p.R255C, p.R264K and p.N299S) in *CASK* were identified in male patients, leading to substitutions in the CaMK domain (Fig. 1, A). All four patients were severely affected by microcephaly, severe developmental delay, intellectual disability and seizures (Table 1). Patient 1 (p.E115K) stands out as he additionally showed PCH and thus had the MICPCH disorder, usually associated with *CASK* loss-of-function mutations. This patient died at a young age. Previously, only three further missense variants in the CaMK domain have been reported, namely p.G178R, p.L209P and p.Y268H (González-Roca *et al*, 2020; Hackett *et al*., 2010; LaConte *et al*, 2019). As it is unclear how N-terminal variants affect protein function, we analysed the functional relevance of the new variants identified here.

**Fig. 1.**
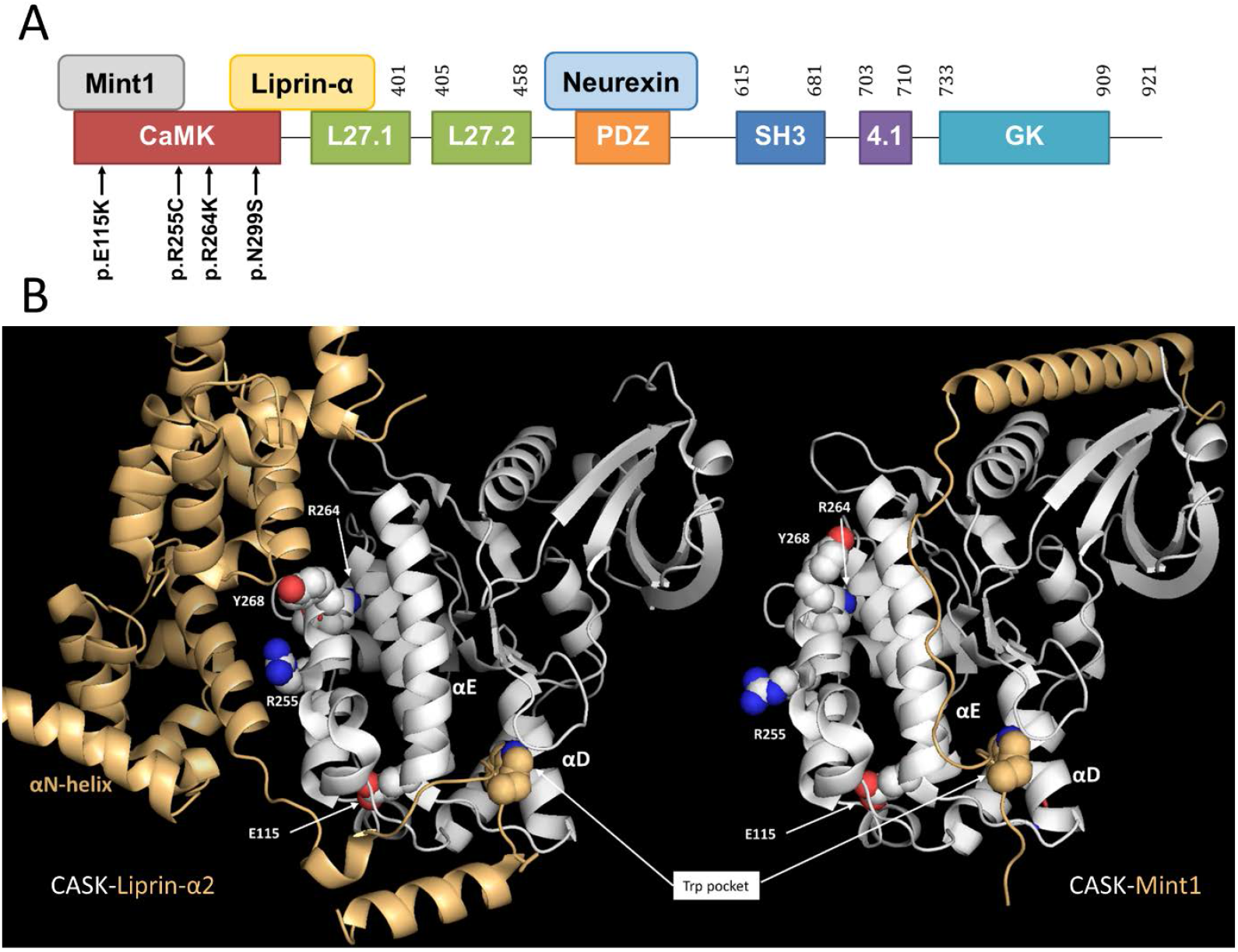
Identification of missense variants affecting the CaMK domain of CASK. **A**. Protein domains of CASK, selected interaction partners and missense variants in the *CASK* gene identified here. Numbering refers to the database entry NP_003679.2. **B**. Comparison of CASK-Liprin-α2 and CASK-Mint1 complexes. The positions of residues analysed here is indicated, with the exception of N299 in the αR1-helix which is occluded by the αD-helix. Note that while the hydrophobic tryptophan pocket is occupied in an identical manner in both complexes, both complexes differ in their secondary binding site, with a cluster of mutants (R255C, R264K, as well as Y268H analysed by (Wei *et al*., 2011)) affecting only the secondary site for Liprin-α2. For Liprin-α2, the position of the αN-helix is also indicated which is pointing away from the SAM domains upon complex formation with CASK. Structures were derived from database entries 3tac in case of Liprin-α2 (Wei *et al*., 2011) and 6kmh in case of Mint1 (Zhang *et al*., 2020) and visualized with Pymol.

**Table 1.**
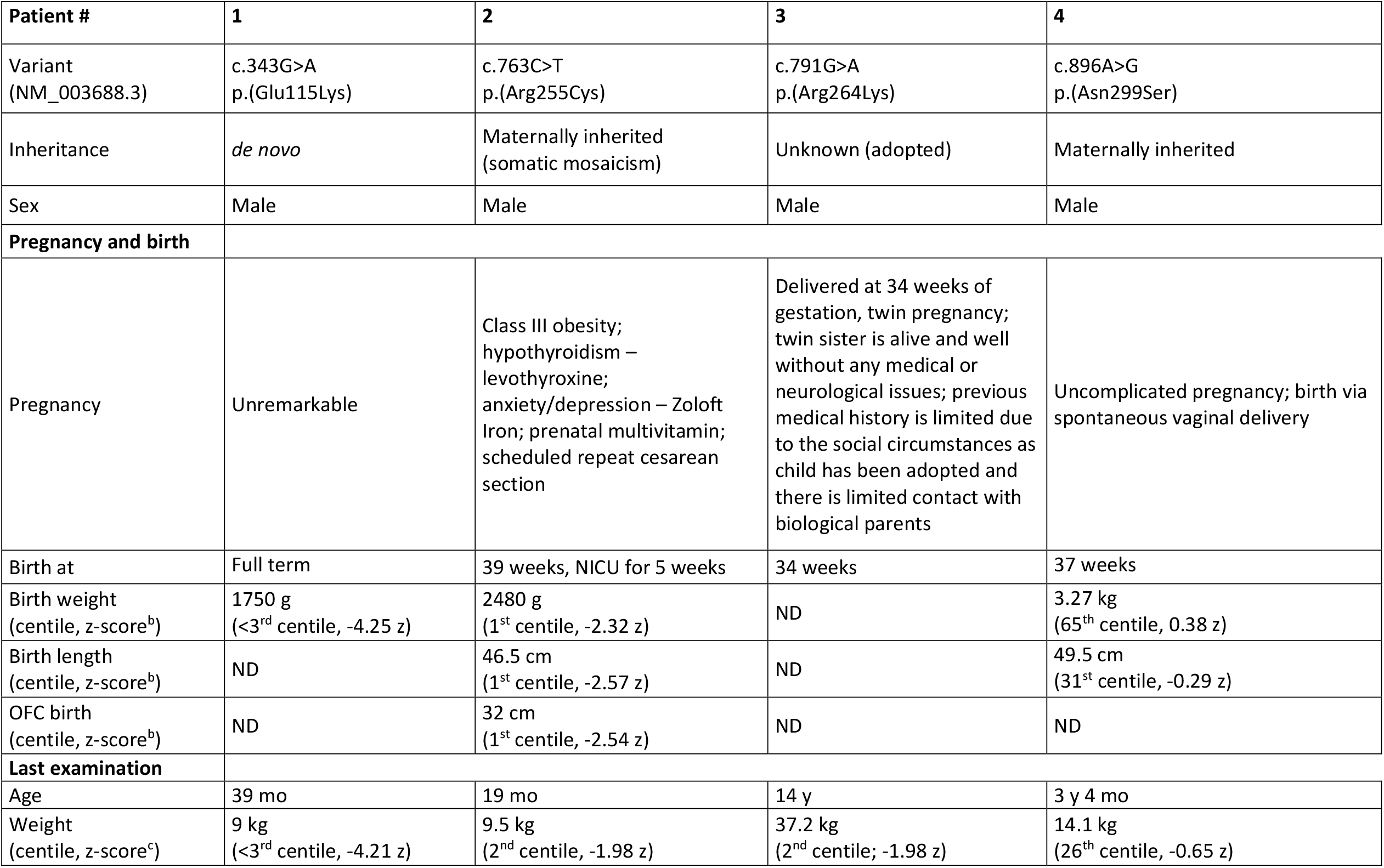
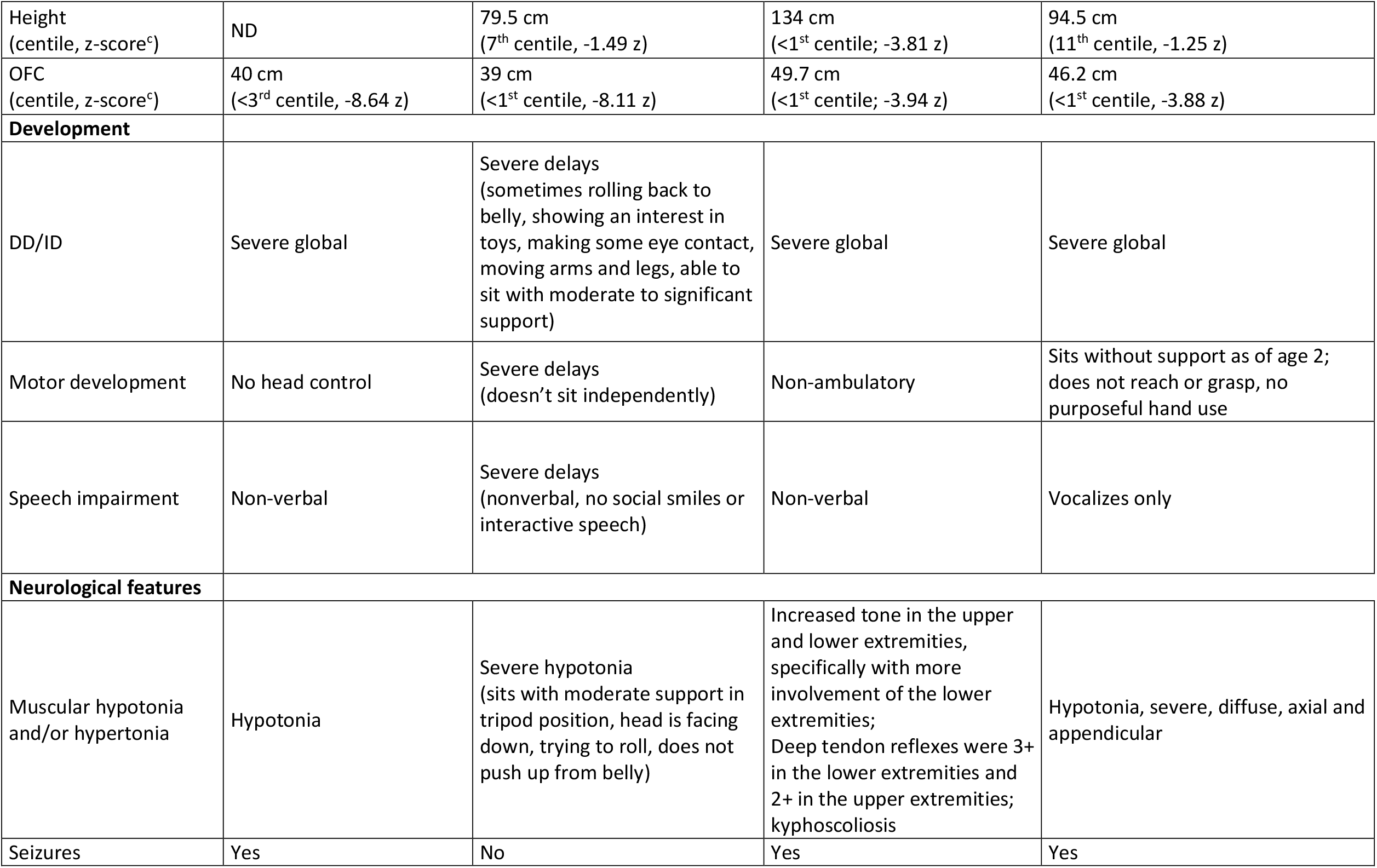
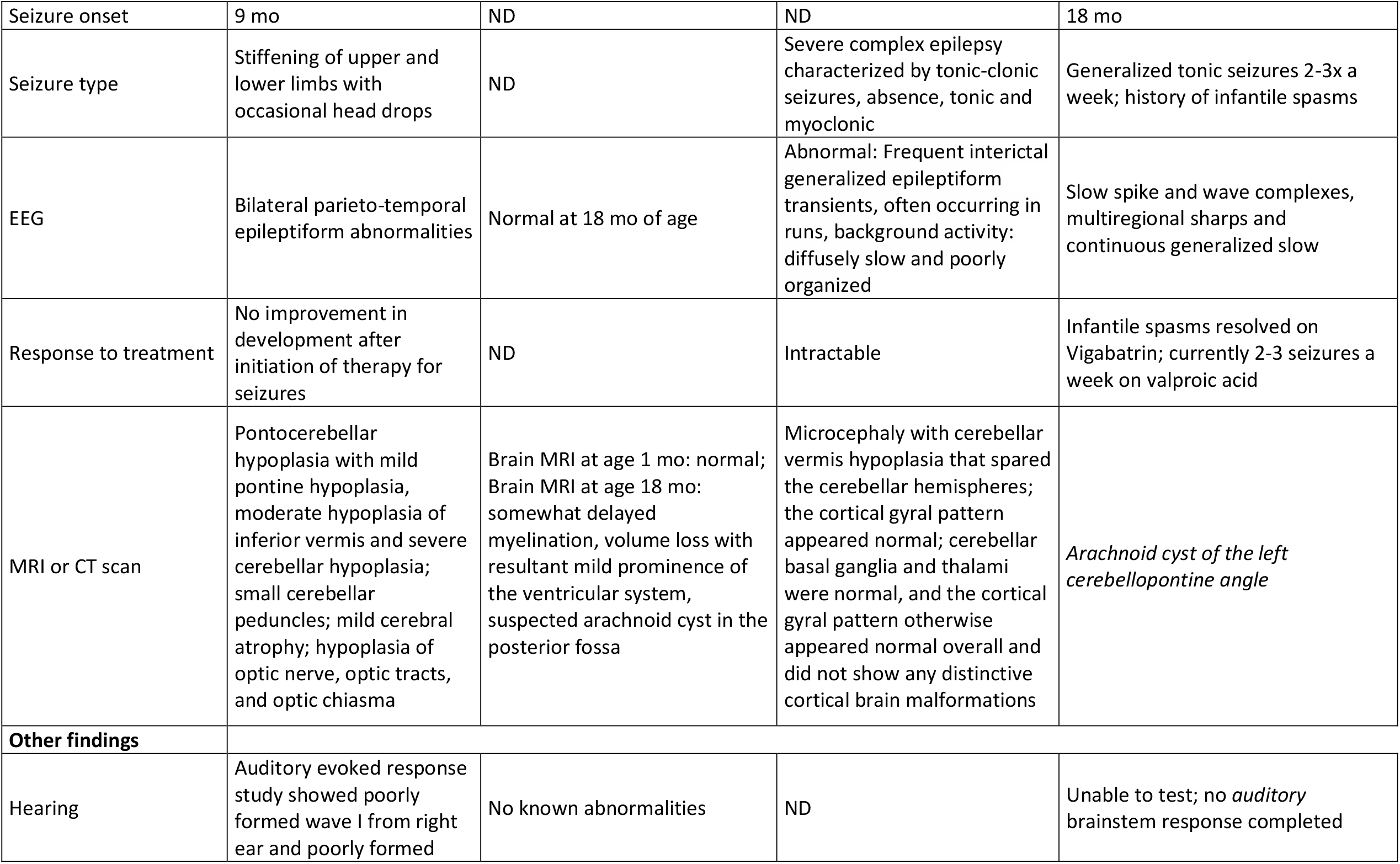
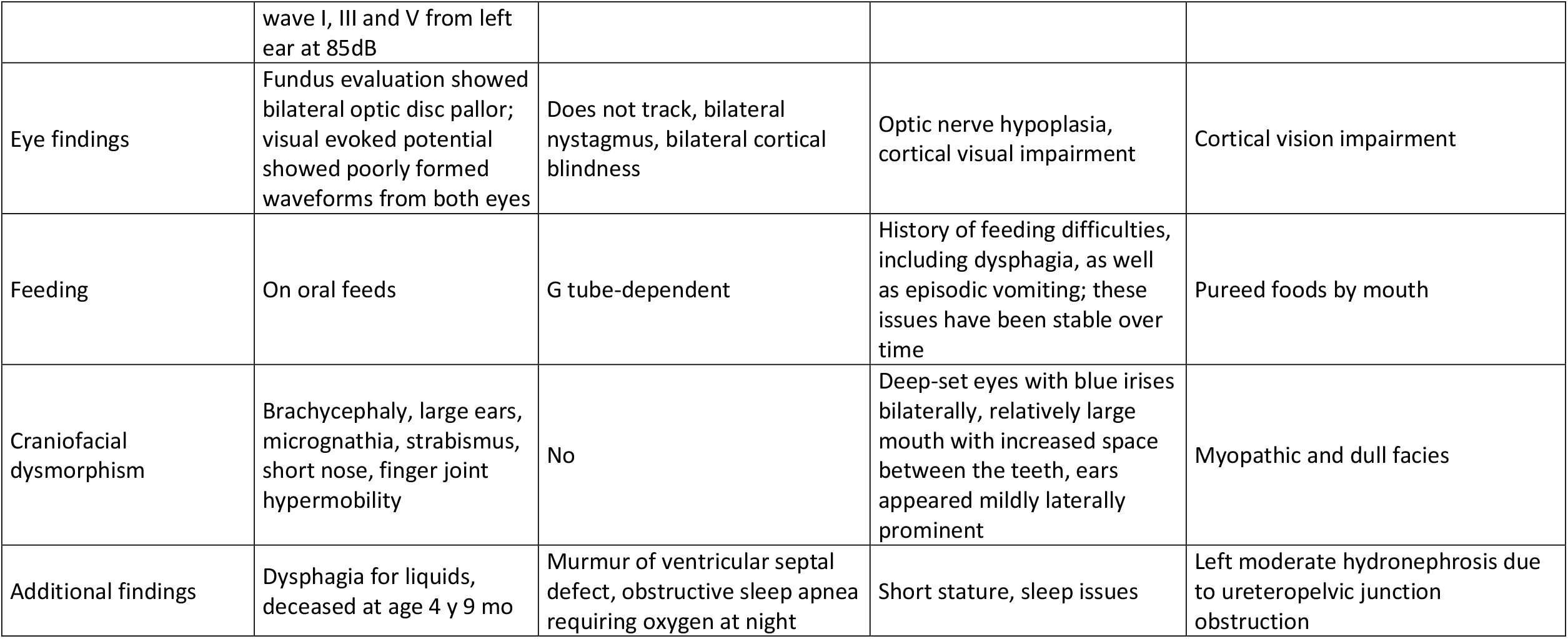
Clinical features in four male patients with a CASK missense variant. ^a^ centiles and z-scores of birth parameters were calculated based on data of (Fenton & Kim, 2013). ^b^ centiles and z-scores were calculated according to Kromeyer-Hauschild, Wabitsch, Kunze et al. Monatsschr Kinderheilkd (2001), 149: 807. https://doi.org/10.1007/s001120170107]. DD, developmental delay; ID, intellectual disability; mo, months; ND, no data; NICU, neonatal intensive care unit; OFC, occipitofrontal head circumference; y, year(s)

| Patient # | 1 | 2 | 3 | 4 |
| --- | --- | --- | --- | --- |
| Variant<br>(NM_003688.3) | c.343G>A<br>p.(Glu115Lys) | c.763C>T<br>p.(Arg255Cys) | c.791G>A<br>p.(Arg264Lys) | c.896A>G<br>p.(Asn299Ser) |
| Inheritance | *de novo* | Maternally inherited<br>(somatic mosaicism) | Unknown (adopted) | Maternally inherited |
| Sex | Male | Male | Male | Male |
| **Pregnancy and birth** |  |  |  |  |
| Pregnancy | Unremarkable | Class III obesity;<br>hypothyroidism –<br>levothyroxine;<br>anxiety/depression – Zoloft<br>Iron; prenatal multivitamin;<br>scheduled repeat cesarean<br>section | Delivered at 34 weeks of<br>gestation, twin pregnancy;<br>twin sister is alive and well<br>without any medical or<br>neurological issues; previous<br>medical history is limited due<br>to the social circumstances as<br>child has been adopted and<br>there is limited contact with<br>biological parents | Uncomplicated pregnancy; birth via<br>spontaneous vaginal delivery |
| Birth at | Full term | 39 weeks, NICU for 5 weeks | 34 weeks | 37 weeks |
| Birth weight<br>(centile, z-score<sup>b</sup>) | 1750 g<br>(<3<sup>rd</sup> centile, -4.25 z) | 2480 g<br>(1<sup>st</sup> centile, -2.32 z) | ND | 3.27 kg<br>(65<sup>th</sup> centile, 0.38 z) |
| Birth length<br>(centile, z-score<sup>b</sup>) | ND | 46.5 cm<br>(1<sup>st</sup> centile, -2.57 z) | ND | 49.5 cm<br>(31<sup>st</sup> centile, -0.29 z) |
| OFC birth<br>(centile, z-score<sup>b</sup>) | ND | 32 cm<br>(1<sup>st</sup> centile, -2.54 z) | ND | ND |
| **Last examination** |  |  |  |  |
| Age | 39 mo | 19 mo | 14 y | 3 y 4 mo |
| Weight<br>(centile, z-score<sup>c</sup>) | 9 kg<br>(<3<sup>rd</sup> centile, -4.21 z) | 9.5 kg<br>(2<sup>nd</sup> centile, -1.98 z) | 37.2 kg<br>(2<sup>nd</sup> centile; -1.98 z) | 14.1 kg<br>(26<sup>th</sup> centile, -0.65 z) |
| Height<br>(centile, z-score<sup>c</sup>) | ND | 79.5 cm<br>(7<sup>th</sup> centile, -1.49 z) | 134 cm<br>(<1<sup>st</sup> centile; -3.81 z) | 94.5 cm<br>(11<sup>th</sup> centile, -1.25 z) |
| OFC<br>(centile, z-score<sup>c</sup>) | 40 cm<br>(<3<sup>rd</sup> centile, -8.64 z) | 39 cm<br>(<1<sup>st</sup> centile, -8.11 z) | 49.7 cm<br>(<1<sup>st</sup> centile; -3.94 z) | 46.2 cm<br>(<1<sup>st</sup> centile, -3.88 z) |
| **Development** |  |  |  |  |
| DD/ID | Severe global | Severe delays<br>(sometimes rolling back to belly, showing an interest in toys, making some eye contact, moving arms and legs, able to sit with moderate to significant support) | Severe global | Severe global |
| Motor development | No head control | Severe delays<br>(doesn't sit independently) | Non-ambulatory | Sits without support as of age 2; does not reach or grasp, no purposeful hand use |
| Speech impairment | Non-verbal | Severe delays<br>(nonverbal, no social smiles or interactive speech) | Non-verbal | Vocalizes only |
| **Neurological features** |  |  |  |  |
| Muscular hypotonia and/or hypertonia | Hypotonia | Severe hypotonia<br>(sits with moderate support in tripod position, head is facing down, trying to roll, does not push up from belly) | Increased tone in the upper and lower extremities, specifically with more involvement of the lower extremities;<br>Deep tendon reflexes were 3+ in the lower extremities and 2+ in the upper extremities;<br>kyphoscoliosis | Hypotonia, severe, diffuse, axial and appendicular |
| Seizures | Yes | No | Yes | Yes |
| Seizure onset | 9 mo | ND | ND | 18 mo |
| Seizure type | Stiffening of upper and lower limbs with occasional head drops | ND | Severe complex epilepsy characterized by tonic-clonic seizures, absence, tonic and myoclonic | Generalized tonic seizures 2-3x a week; history of infantile spasms |
| EEG | Bilateral parieto-temporal epileptiform abnormalities | Normal at 18 mo of age | Abnormal: Frequent interictal generalized epileptiform transients, often occurring in runs, background activity: diffusely slow and poorly organized | Slow spike and wave complexes, multiregional sharps and continuous generalized slow |
| Response to treatment | No improvement in development after initiation of therapy for seizures | ND | Intractable | Infantile spasms resolved on Vigabatrin; currently 2-3 seizures a week on valproic acid |
| MRI or CT scan | Pontocerebellar hypoplasia with mild pontine hypoplasia, moderate hypoplasia of inferior vermis and severe cerebellar hypoplasia; small cerebellar peduncles; mild cerebral atrophy; hypoplasia of optic nerve, optic tracts, and optic chiasma | Brain MRI at age 1 mo: normal; Brain MRI at age 18 mo: somewhat delayed myelination, volume loss with resultant mild prominence of the ventricular system, suspected arachnoid cyst in the posterior fossa | Microcephaly with cerebellar vermis hypoplasia that spared the cerebellar hemispheres; the cortical gyral pattern appeared normal; cerebellar basal ganglia and thalami were normal, and the cortical gyral pattern otherwise appeared normal overall and did not show any distinctive cortical brain malformations | *Arachnoid cyst of the left cerebellopontine angle* |
| **Other findings** |  |  |  |  |
| Hearing | Auditory evoked response study showed poorly formed wave I from right ear and poorly formed wave I, III and V from left ear at 85dB | No known abnormalities | ND | Unable to test; no *auditory* brainstem response completed |
| Eye findings | Fundus evaluation showed bilateral optic disc pallor; visual evoked potential showed poorly formed waveforms from both eyes | Does not track, bilateral nystagmus, bilateral cortical blindness | Optic nerve hypoplasia, cortical visual impairment | Cortical vision impairment |
| Feeding | On oral feeds | G tube-dependent | History of feeding difficulties, including dysphagia, as well as episodic vomiting; these issues have been stable over time | Pureed foods by mouth |
| Craniofacial dysmorphism | Brachycephaly, large ears, micrognathia, strabismus, short nose, finger joint hypermobility | No | Deep-set eyes with blue irises bilaterally, relatively large mouth with increased space between the teeth, ears appeared mildly laterally prominent | Myopathic and dull facies |
| Additional findings | Dysphagia for liquids, deceased at age 4 y 9 mo | Murmur of ventricular septal defect, obstructive sleep apnea requiring oxygen at night | Short stature, sleep issues | Left moderate hydronephrosis due to ureteropelvic junction obstruction |

To assess the potential of the CASK mutants to disrupt specific functions of CASK, we looked at 3D crystal structures of the CASK CaMK domain in complexes with Mint1 and Liprin-α2, two prominent presynaptic partners of CASK. Both Mint1 and Liprin-α2 use the insertion of a tryptophan side chain which is part of a conserved Ile/Val-Trp-Val sequence (Trp981 in Liprin-α2) into a deep hydrophobic pocket in CASK (Wei *et al*., 2011; Wu *et al*, 2020). This pocket is formed between the αD and αE helices of the kinase domain (Fig. 1, B). As the αD helix is intimately connected to the αR1 helix in CASK, it is conceivable that the N299S variant in αR1 might alter the hydrophobic pocket. The E115K variant in αE is likely to affect the position of αE, thereby changing the size of the pocket. Liprin-α2 requires a second interface for high affinity binding, which is formed by its SAM2 domain. For CASK, this involves part of the CaMK surface containing residues R255, R264 and Y268. The substitutions R255C and R264K (as well as Y268H, analysed by (Wei *et al*., 2011)) therefore are likely to affect the second interface between CASK and the SAM2 domain of Liprin-α2.

### Binding to Mint1 and Neurexin is not affected whereas binding to Veli and ATP is slightly altered by missense variants affecting the CASK CaMK domain

CASK forms an evolutionarily conserved trimeric complex with Mint1 and Veli proteins (Butz *et al*., 1998). The interaction between CASK and Mint1 is mediated by the CaMK domain of CASK (Wu *et al*., 2020; Zhang *et al*, 2020). We tested whether the variants altered binding of CASK to Mint1 or Veli. HEK293T cells were cotransfected with plasmids coding for mRFP-tagged CASK variants (or mRFP alone as negative control) and GFP-tagged Mint1. Veli proteins are highly expressed endogenously in HEK293T cells and were visualized in two bands at about 26 and 30 kDa, corresponding to Veli1 at 30 kDa and Veli2/3 at 26 kDa. Upon cell lysis and immunoprecipitation of mRFP-containing proteins, we found that all CASK mutant variants interact consistently well with Mint1 and Veli proteins. Veli2/3 were more prominent in the IP sample compared to Veli1 (Fig. 2, A). The E115K variant co-precipitated slightly more Veli proteins than CASK-WT and the remaining mutants (Fig. 2, A-C).

**Fig. 2.**
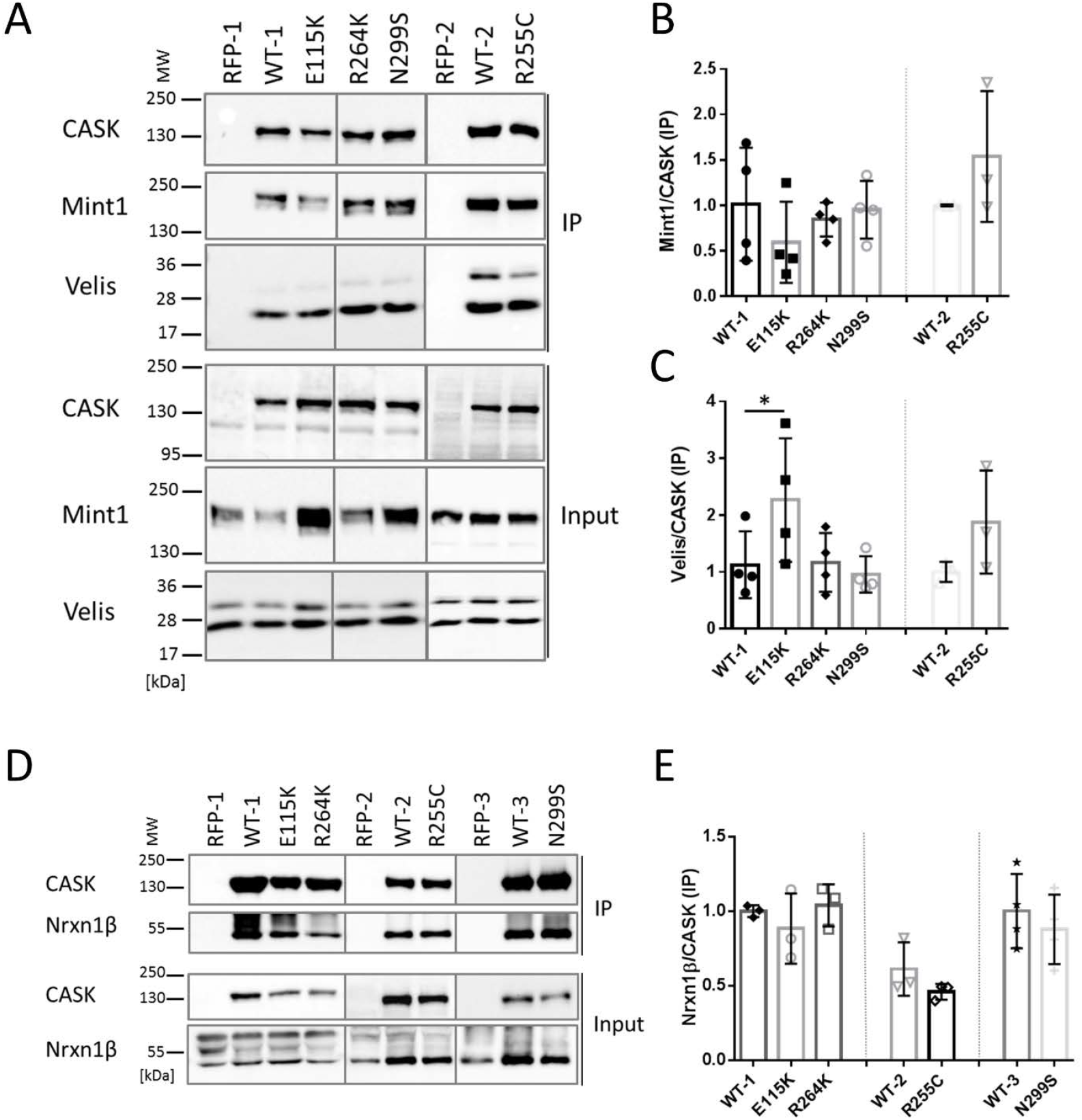
Substitutions in the CaMK domain of CASK do not alter interactions with Mint1 and Neurexin whereas binding to Veli is slightly increased for the E115K variant. **A**. mRFP-tagged CASK-WT and mutants, or mRFP alone were coexpressed with Mint1 in HEK293T cells. mRFP-tagged proteins were immunoprecipitated from cell lysates, and inputs (IN) and precipitates (IP) were analysed by western blotting using antibodies against GFP- and mRFP-tags, as well as anti-Veli. **B, C**. Quantitative analysis of results shown in B. Coprecipitation efficiency was determined as the ratio of Mint1 (B) or Veli (C) IP signal, divided by the CASK IP signal. **D, E**. mRFP-tagged CASK variants were coexpressed with HA-tagged Neurexin, and mRFP-tagged proteins were immunoprecipitated as in B. Quantitation in E shows that Neurexin-1β binding was not affected by CASK variants. The mean ± *SD* is shown with each data point representing an independent transfection experiment. Significance was determined by one-way ANOVA with *post hoc* Dunnett’s multiple comparisons test or two-tailed Student’s *t*-test; *, p≤0.05; n=3-4.

The C-terminal PDZ ligand of Neurexins binds to the PDZ domain of CASK (Hata *et al*., 1996). Structural work has shown that the full PSG superdomain is necessary for high affinity interaction (Li *et al*., 2014) and oligomerization (Pan *et al*., 2021). Importantly, the C-terminus of Neurexin is the only known *in vivo* substrate for the CASK CaM kinase activity (Mukherjee *et al*, 2010; Mukherjee *et al*., 2008), suggesting the possibility that alterations in the CaMK domain might affect the interaction between both proteins. To test this, HEK293T cells were transiently transfected with plasmids coding for mRFP-CASK and HA-Neurexin-1β. As before, mRFP-tagged CASK variants (or mRFP as control) were immunoprecipitated from cell lysates. Amounts of CASK and coprecipitated Neurexin were analysed by western blot and quantified (Fig. 2, D and E). With all CASK variants, Neurexin-1β was coimmunoprecipitated efficiently and no differences with respect to the CASK-Neurexin interaction were detected.

For measurement of ATP binding to the active site of the kinase domain we employed the fluorescent ATP analog TNP-ATP. WT and mutant CASK kinase domains were expressed as His_6_-tag-SUMO fusion proteins in bacteria and purified. Binding of TNP-ATP to both WT and mutant CASK kinase domains could be verified by a shift of the fluorescence spectrum of TNP-ATP to lower wavelengths, and an increase in fluorescence intensity (Fig. 3, A). TNP-ATP binding was reduced in the presence of Mg^2+^, in keeping with the designation of CASK as an atypical, Mg^2+^-sensitive instead of Mg^2+^-dependent, kinase (Mukherjee *et al*., 2008). ATP binding to all CASK variants proved to be Mg^2+^ sensitive. By calculating the ratio of fluorescence in the absence and in the presence of Mg^2+^ as the “Mg^2+^ sensitivity” we determined that the R264K and N299S variants of CASK were significantly more sensitive to Mg^2+^ than the WT protein and the other mutants, E115K and R255C (Fig. 3, B and C).

**Fig. 3.**
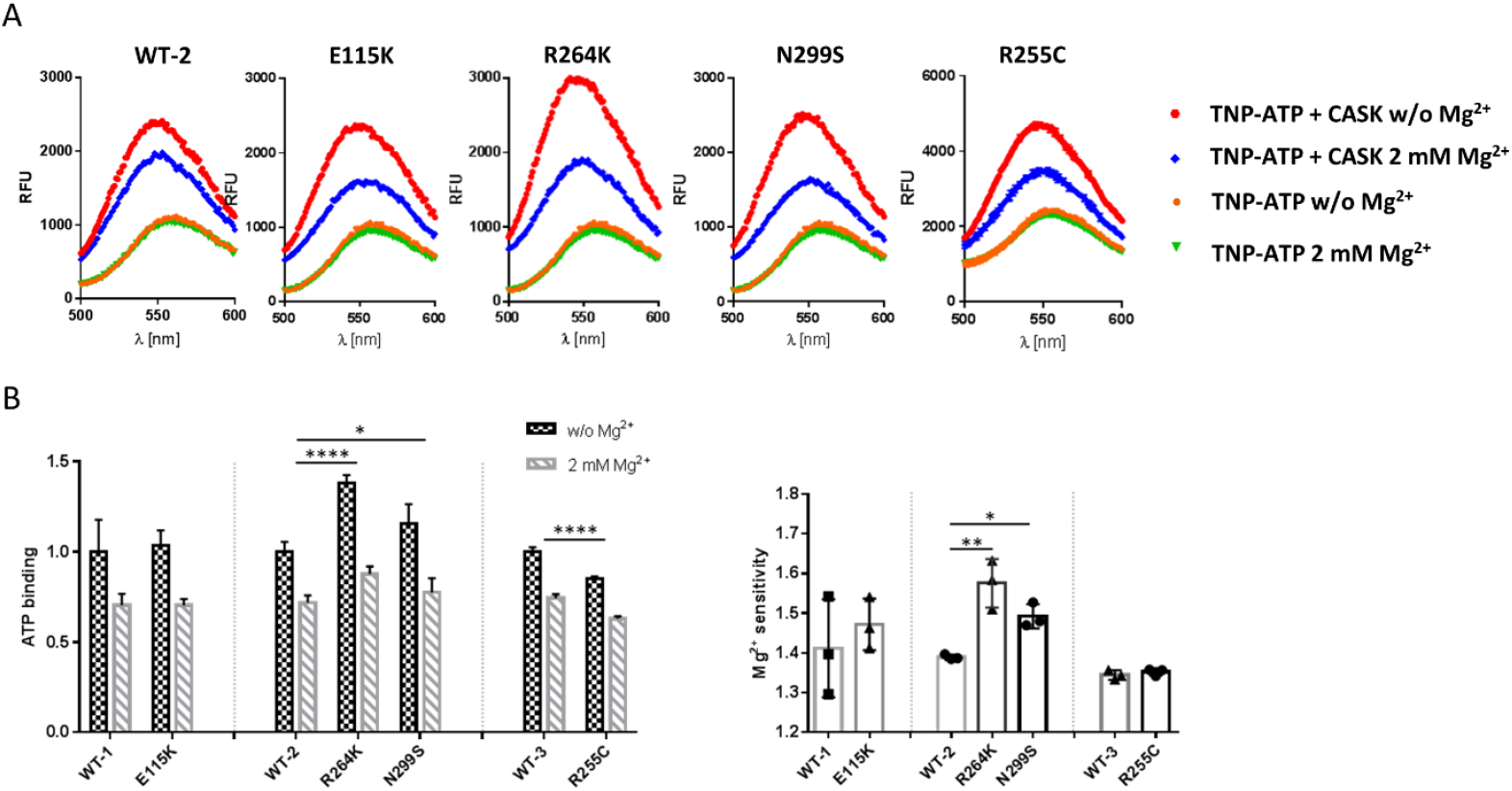
Mg^2+^-sensitivity of ATP binding is slightly altered by R264K and N299S variants of CASK. SUMO-tagged fusion proteins of the CaMK domain of CASK were purified and incubated with the fluorescent ATP analog TNP-ATP in the absence or presence of 2 mM Mg^2 +^ and fluorescence emission spectra were recorded from 500 nm to 600 nm. **B, C**. Quantification of the maxima of fluorescence signal in the absence and presence of Mg^2+^ (B), and of the ratio between the two values, defined as the “Mg^2+^ sensitivity” (C). Significance was determined by (B) two-way ANOVA with Sidak’s multiple comparison test or (C) one-way ANOVA with Dunnett’s multiple comparisons test or two-tailed Student’s *t*-test; *, p≤0.05, **, p≤0.01; ****, p≤0.0001; n=3. Mean ± *SD* is shown. In C, each data point represents an independent fusion protein purification.

### Reduced binding to Liprin-α2

To analyse whether binding to the active zone component Liprin-α2 (LaConte *et al*., 2016; Olsen *et al*, 2005) is affected, HEK293T cells were transfected with plasmids coding for either a CASK variant or the empty vector control together with a construct coding for HA-tagged Liprin-α2. Upon immunoprecipitation of mRFP-tagged proteins, we found that all four tested CaMK mutants showed a significant reduction in interaction with Liprin-α2 compared to CASK-WT (Fig. 4, A and B). We repeated this assay in a different format, using an immobilized SUMO fusion protein of the three C-terminal SAM domains of Liprin-α2, which contain the CASK binding loop (Wei *et al*., 2011), in a pulldown assay from HEK293T cells expressing the different mRFP-tagged CASK variants. Here, we similarly observed differences between CASK-WT and the four mutants, as significantly reduced binding was detected for E115K, R255C and R264K. The reduction in binding for the N299S variant was not significant after three repeats (Fig. 4, C and D).

**Fig. 4.**
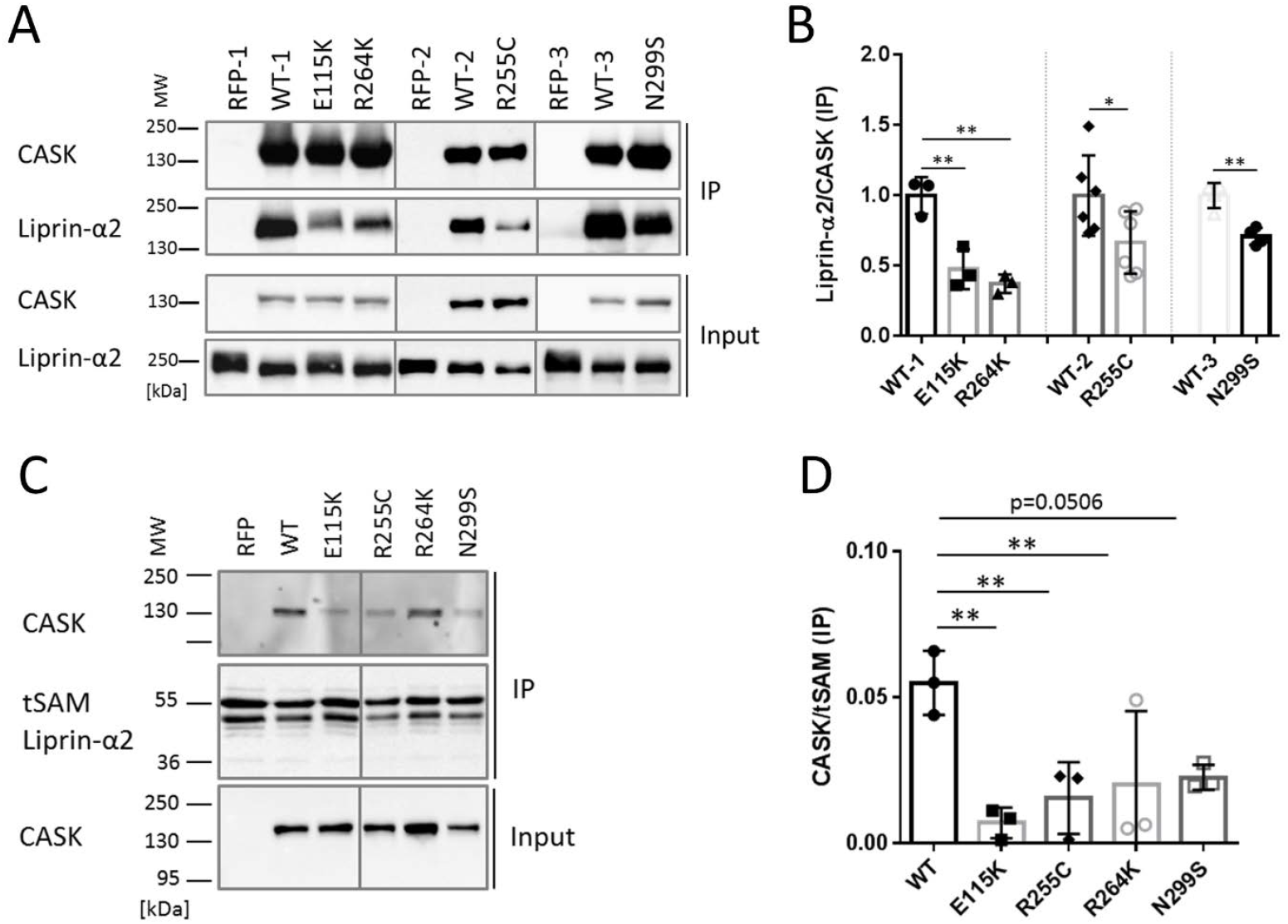
All four *CASK* variants interfere with binding to Liprin-α2. **A**. HA-tagged Liprin-α2 was coexpressed with mRFP or mRFP-tagged CASK variants. mRFP-containing proteins were immunoprecipitated from cell lysates, and input and precipitate samples were analysed by western blotting using epitope-specific antibodies. **B**. Quantification of the data shown in A as mean ± *SD*. Precipitation efficiency is in each case quantified as the ratio of precipitated HA-Liprin signal divided by precipitated mRFP-CASK signal. **C**. A His6-Sumo fusion protein of the region encompassing the C-terminal three SAM domains of Liprin-α2 (tSAM) was isolated from bacteria using Ni-NTA agarose and left on agarose beads for use in a pulldown assay. Beads were incubated with lysates from cells expressing mRFP alone or mRFP-tagged CASK variants. After washing, input and precipitate samples were analysed by western blotting using the antibodies indicated. D. Quantification of the data in C shown as mean ± *SD*. Interaction was quantified as the ratio of CASK signal in precipitates, divided by the signal of the tSAM domains. Statistics in (B) and (D) done with two-tailed Student’s *t*-test or one-way ANOVA followed by Dunnett’s multiple comparison test, respectively; *, p≤0.05, **, p≤0.01; n=3-6.

One might ask why binding of CASK to Liprin is reduced by the variants whereas binding to Mint1 is not. Mint1 also uses the hydrophobic pocket of CASK for insertion of its Val-Trp-Val sequence; however, Mint1 uses a second binding interface distinct from that of Liprin-α2. This involves a stretch of α-helix which makes an extensive contact to the N-lobe of the CASK CaMK domain (Fig. 1, B) (LaConte *et al*., 2016; Wu *et al*., 2020), allowing for a much higher affinity of the Mint1-CASK interaction (Wu *et al*., 2020). We think that Mint1 is not affected by the variants because (a) R255 and R264 are in the second interface for Liprin-α2 but not for Mint1 (Fig. 1, B); and (b) because the second interface for Mint1 provides more strength to the interaction. Thus, disruption of the hydrophobic pocket for the Trp side chain can be partially compensated by the second interface of Mint1, but not of Liprin-α2.

We focused further on the interaction between CASK and Liprin-α2. In Fig. 4, A; and more pronounced in Fig. 5, A; we noted that the Liprin-α2 band in western blot analysis smeared and was shifted to higher molecular weights when coexpressed with mRFP from the empty vector, but not when coexpressed with CASK-WT. Upward smearing of Liprin-α2 was also observed when CASK-E115K was coexpressed; R264K and N299S showed a somewhat intermediary effect (Fig. 5, A). As we suspected a phosphorylation event, lysates of cells expressing mRFP-CASK or mRFP alone together with Liprin-α2 were treated with FastAP phosphatase. This treatment eliminated the shift to higher molecular weight (Fig. 5, B). Coexpression of CASK-WT with Liprin-α2 led to higher electrophoretic mobility of Liprin-α2, which was similar to that of the phosphatase treated samples in the absence of CASK. No further increase in mobility was observed when the CASK+Liprin-α2 samples were treated with phosphatase (Fig. 5, B). As a conclusion, Liprin-α2 is phosphorylated in 293T cells in the absence of CASK; coexpression of CASK-WT but not CASK-E115K interfered with this phosphorylation event in Liprin-α2.

**Fig. 5.**
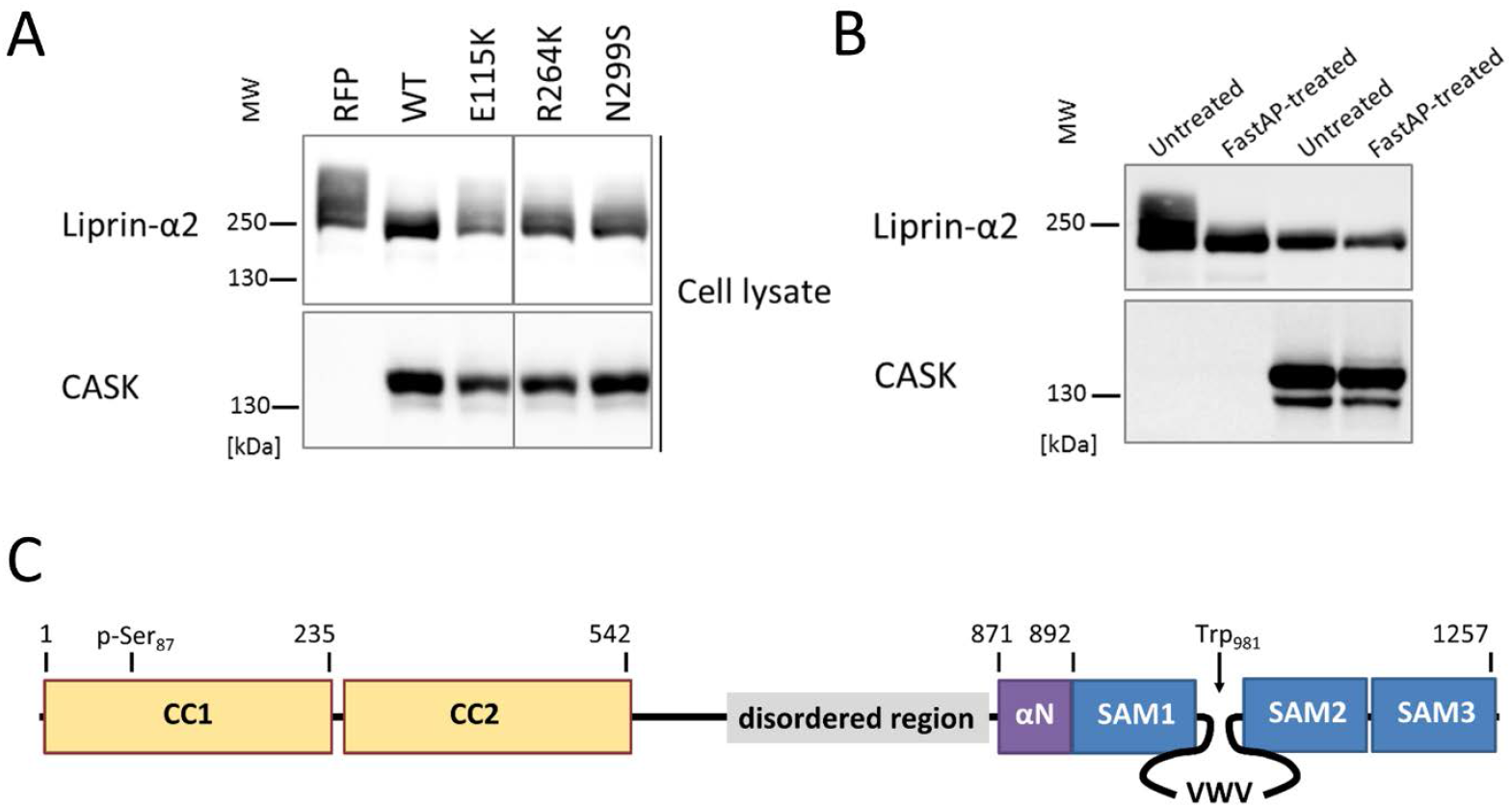
Interaction with CASK alters the phosphorylation status of Liprin-α2. **A**. Lysates from cells coexpressing HA-tagged Liprin-α2 with mRFP or mRFP-tagged variants of CASK were analysed by SDS-PAGE on an 8 % gel, followed by western blotting. Note the upward smear of the Liprin-α2 specific band, which is abolished by coexpression with CASK-WT but not by CASK mutants. **B**. Lysates from cells coexpressing HA-tagged Liprin-α2 with mRFP or mRFP-tagged CASK_WT were treated with or without the FastAP alkaline phosphatase, followed by analysis by SDS-PAGE on an 8 % gel and western blotting. **C**. Domain structure of Liprin-α2; the positions of Ser87 which is most strongly phosphorylated in the absence of CASK, and of Trp981, which constitutes a major binding interface for CASK, are indicated. CC, coiled coil.

We determined phosphorylation sites by mass spectroscopic analysis of immunoprecipitated Liprin-α2, isolated from cells coexpressing mRFP control vector, or coexpressing mRFP-tagged CASK. Numerous phosphorylated sites were detected in the N-terminal coiled-coil regions (i.e. S73, S87, S257, S260, S263) and the intervening IDR (S552 and S673). Remarkably, phosphorylation of S87 in coiled-coil region 1 was decreased when CASK was coexpressed in three independent repeats (Fig. 5, C, Table 2).

**Table 2.**
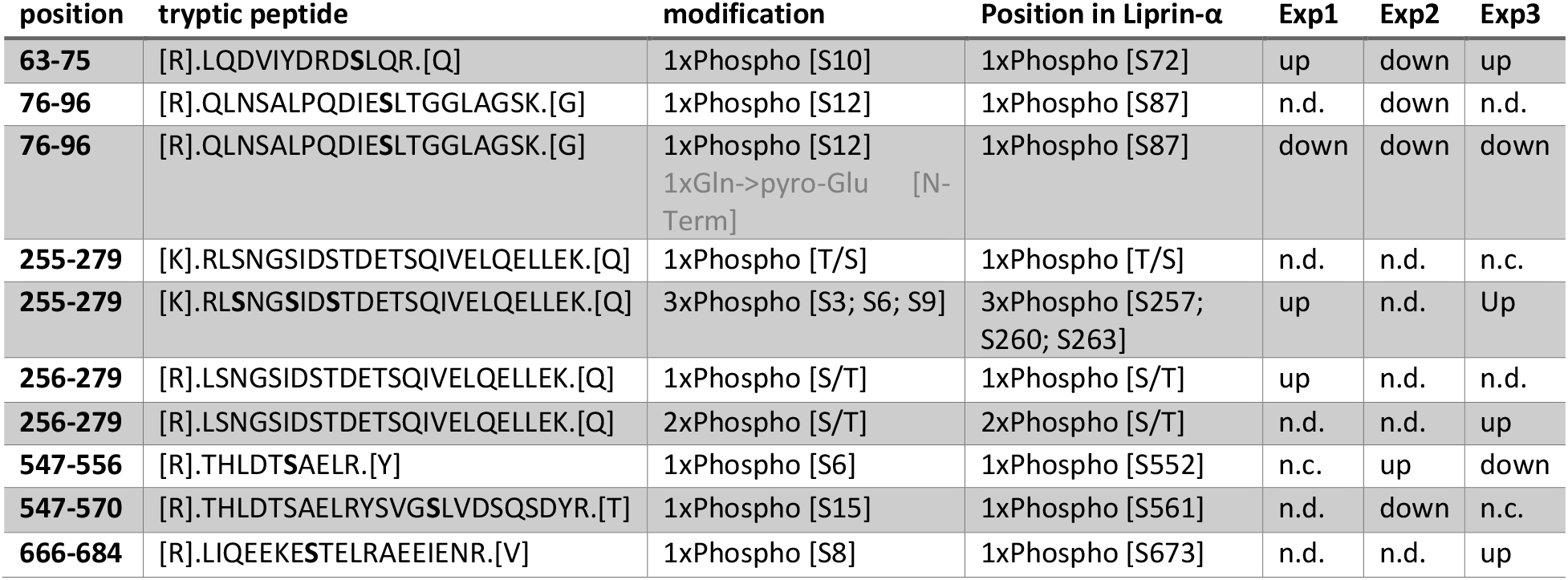
CASK coexpression alters phosphorylation of Liprin-α2. 293T cells expressing Liprin-α2 alone or in combination with CASK-WT were lysed, and HA-tagged Liprin was immunoprecipitated using anti-HA magnetic beads. Purified samples were analysed by tryptic digestion, followed by mass spectroscopy. Phosphorylated peptides were identified and quantified in three independent experiments (Exp1-3). “up” and “down” denotes increases and decreases in peptide intensity upon coexpression with CASK, respectively; n.c., no change; n.d., not detected.

Liprin-α proteins from various species form condensates in cells through a process termed liquid-liquid phase separation (LLPS) (Emperador-Melero *et al*., 2021; Liang *et al*., 2021; McDonald *et al*, 2020; Xie *et al*., 2021). During LLPS, proteins form dynamic, non-membrane surrounded compartments through demixing from the diffuse state out of the cytosol (Bracha *et al*, 2019). The homolog of Liprin-α in *C. elegans* (SYD-2) exhibits liquid-liquid phase separation in early stages of synapse development (McDonald *et al*., 2020). Upon microscopic analysis of Liprin-α2 expressing 293T cells, we observed large spherical condensates of Liprin-α2 which appeared to be LLPS-like events such as those observed by (Emperador-Melero *et al*., 2021) (Fig. 6). Coexpression of Liprin-α2 with CASK-WT resulted in a diffuse cytosolic localization for both proteins in the majority of analysed cells. This CASK-dependent change of intracellular localization from bigger condensates to a cytosolic diffuse localization was also observed upon coexpression with CASK variants R255C, R264K and N299S (Fig. 6). In a striking contrast, the localization of Liprin-α2 resembled that observed in the absence of overexpressed CASK when the CASK-E115K variant was coexpressed. CASK-E115K colocalized with Liprin-α2 in the large condensates that were formed (Fig. 6). Thus, CASK was able to negatively regulate condensate formation, and CASK-E115K failed to do so. To further investigate this phenomenon, we performed a time resolved series of experiments. Here, cells were first transfected with the Liprin-α2 construct to allow for formation of spherical droplets. On the next day, cells were transfected again with CASK expression vectors. Here we observed that CASK-WT was indeed able to “dissolve” preformed condensates of Liprin-α2 (Fig. 7).

**Fig. 6.**
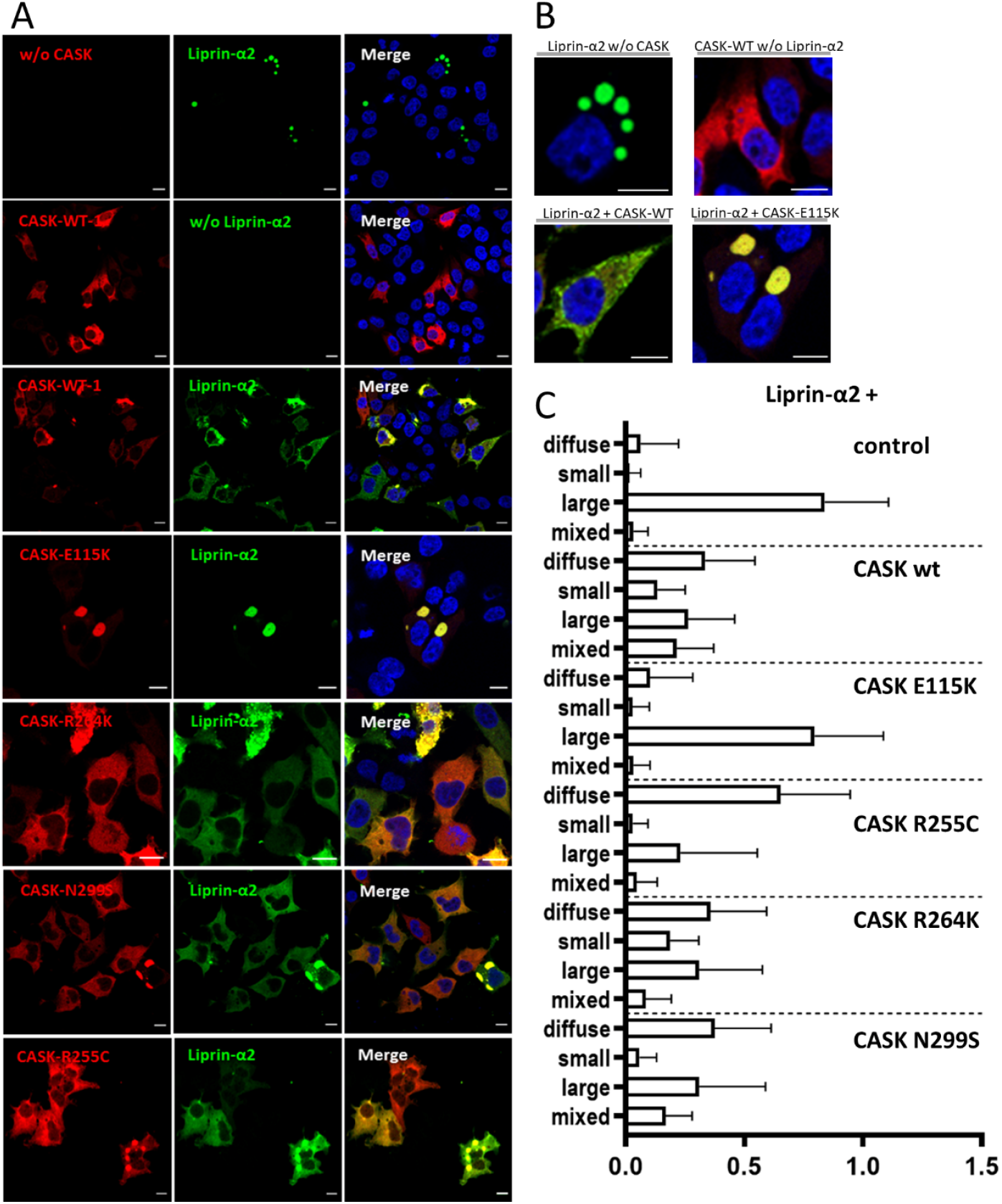
Interaction with CASK interferes with formation of spherical condensates by Liprin-α2 in HEK293T cells. **A**. 293T cells expressing GFP-Liprin-α2, mRFP-tagged CASK, or combinations of both proteins were fixed and imaged by confocal microscopy. Images of DAPI-stained nuclei were included in merged pictures. **B**. Enlargements of cells expressing GFP-Liprin-α2 or mRFP-CASK-WT alone, or combinations of GFP-Liprin-α2 with WT or E115K-mutant CASK. **C**. Quantification of data shown in A. Five microscopic fields of view were evaluated per independent experiment. All transfected cells were grouped by the following criteria: diffuse; small clusters, large clusters, mixed small and large clusters. Shown is the percentage of cells in each group based on the total number of transfected and analysed cells. For each condition, three independent experiments with a total of more than 90 cells were evaluated.

**Figure 7.**
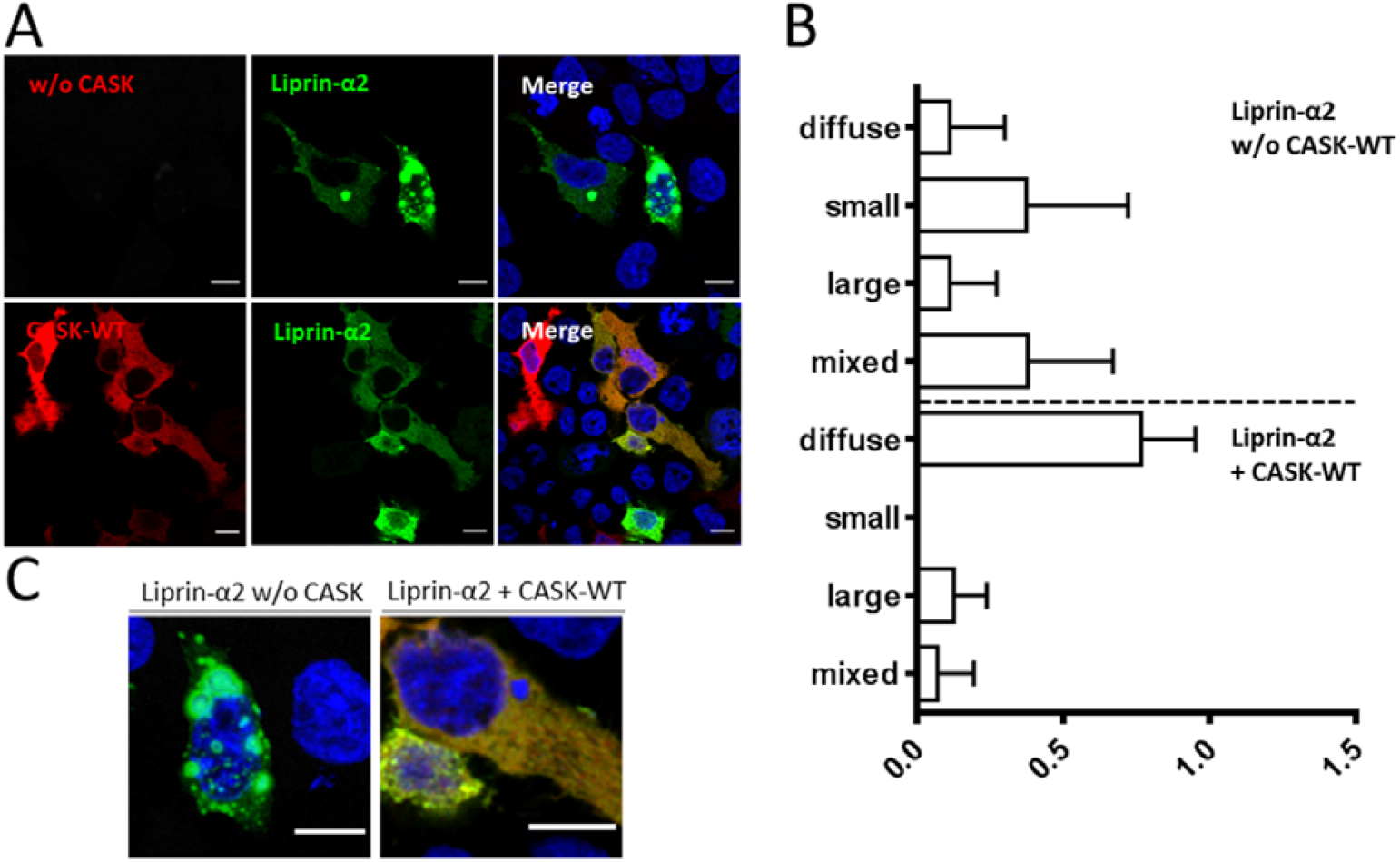
CASK-WT negatively regulates preformed LLPS-like condensates of Liprin-α2. **A**. HEK293T cells were transfected with an expression vector for GFP-tagged Liprin-α2. After two days, cells were fixed and processed for confocal microscopy (upper panels). Alternatively, cells were retransfected with mRFP-CASK after one day, and fixed on the second day. **B**. Quantitative analysis of the data shown in A; categorization and counting of cells was performed as described in Fig. 6. **C**. Enlargement of typical cells shown in A.

In the next set of experiments we asked how CASK and Liprin-α2 affected their mutual localization in primary cultured hippocampal neurons. Here, Liprin-α2 expressed alone was found in condensates throughout the cell bodies, dendrites and axons of transfected neurons. Upon coexpression of CASK-WT, this situation changed as both Liprin-α2 and CASK were localized in a diffuse pattern throughout the cell. Again, the CASK-E115K variant failed to alter the distribution of Liprin-α2, as before in 293T cells (Fig. 8). Closer inspection of axons showed that CASK-WT and Liprin-α2 frequenly colocalized at vGlut1-positive, presumably presynaptic terminals. In contrast, the large LLPS-like condensates observed in cells coexpressing CASK-E115K and Liprin-α2, were found along the axon but were not colocalized with vGlut1, indicating that these are not functional synaptic sites (Fig. 9).

**Fig. 8.**
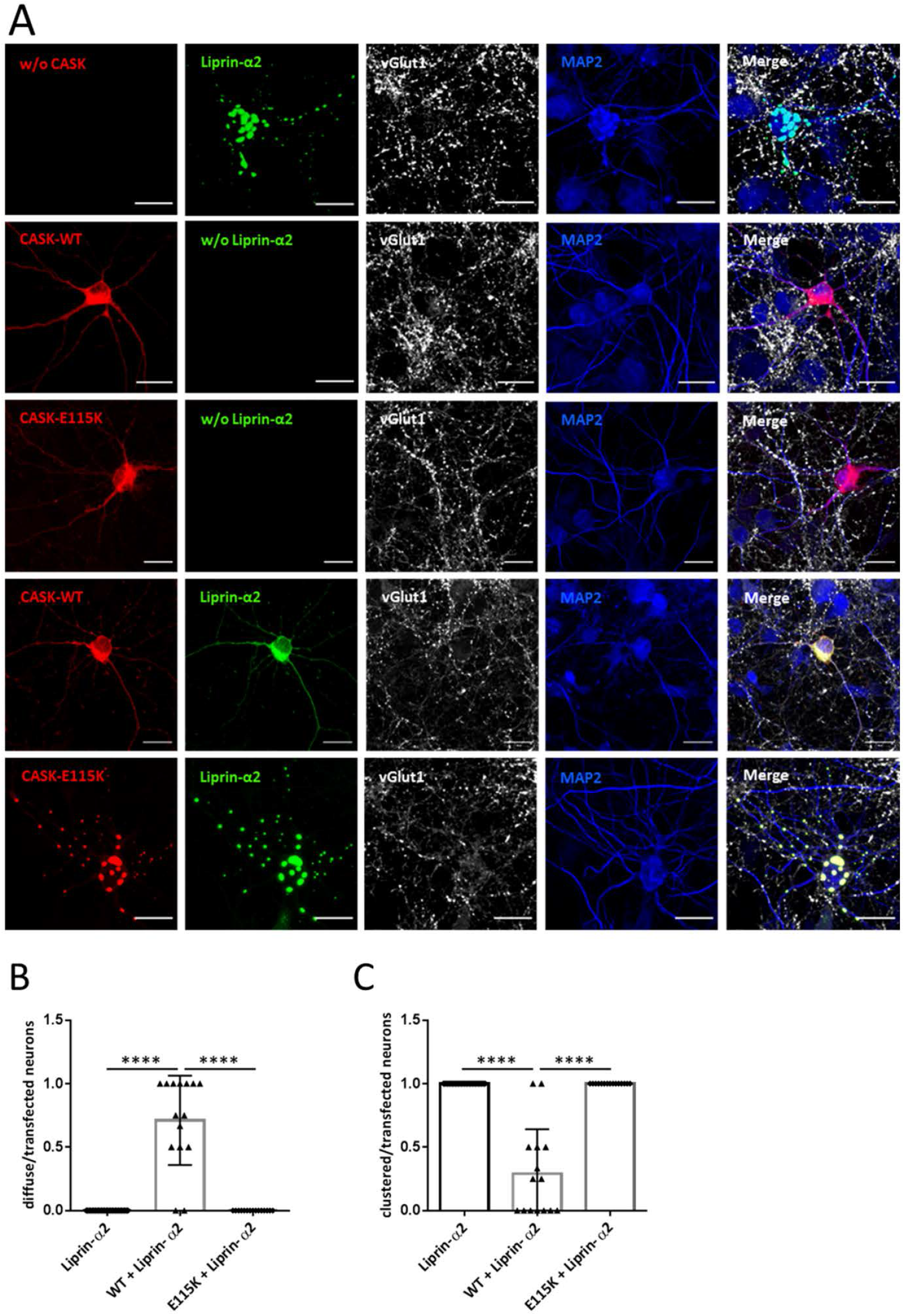
Interaction with CASK interferes with formation of condensates by Liprin-α2 in hippocampal neurons. **A**. Hippocampal neurons transfected with constructs coding for either mRFP-CASK or GFP-Liprin-α2, or both proteins in combination, were fixed and stained for the expressed proteins, as well as vGlut1 as a presynaptic marker and MAP2 as a dendrite marker. **B, C**. Quantification of the data shown in A. **B**. Proportion of neurons in which Liprin-α2 was localized diffusely in the cytoplasm of soma and neurites, normalized to the number of transfected cells per image. **C**. Number of cells in which Liprin-α2 showed a localization in clusters in cell soma and along the neurites. 15 images per condition were analysed with up to five transfected neurons present and the mean ± *SD* is shown with each data point representing one analysed picture. Statistical differences were calculated using an ordinary one-way ANOVA with Tukey’s test; ****, p≤0.0001.

**Fig. 9.**
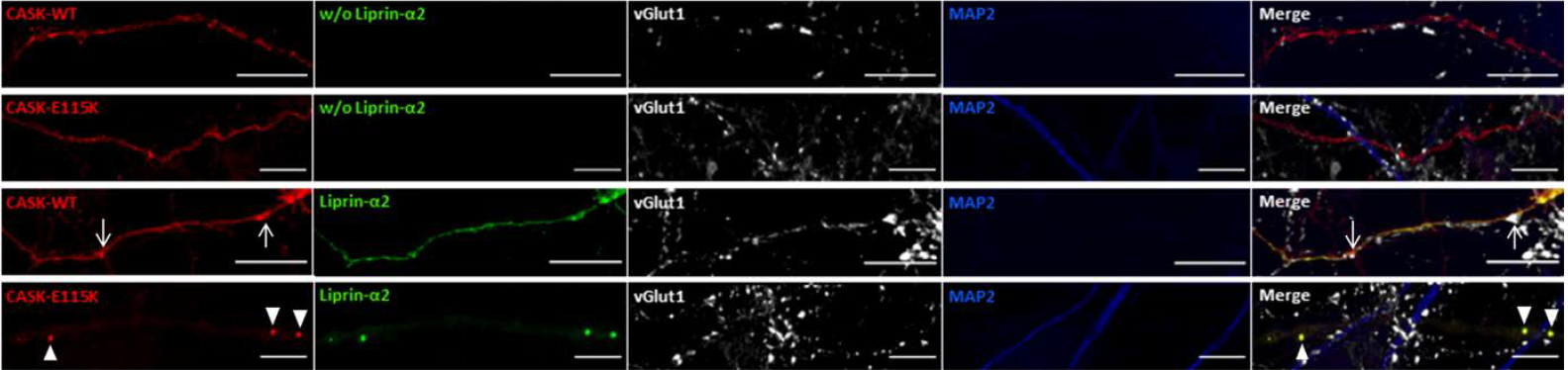
Liprin-α2 condensate-like axonal puncta are not synaptic. Enlargements of axonal segments of neurons shown in Fig. 6. Axons were identified by the absence of MAP2 staining. Arrows point to locations of presynaptic sites identified by vGlut1 staining; arrowheads in lower panels point to CASK-Liprin droplets which are devoid of a presynaptic vGlut1 cluster, and are therefore considered as non-synaptic.

MAGUKs like CASK or PSD-95 are known to oligomerize through their C-terminal PSG tandem domains (McGee *et al*, 2001; Pan *et al*., 2021; Rademacher *et al*, 2019). We analysed the relation between CASK oligomerization and condensation of Liprin-α2, by using a split-YFP fluorescent complementation assay. Two CASK cDNA variants were expressed, carrying either the N-terminal, or C-terminal half of YFP. We have shown before that, upon assembly of CASK oligomers, this leads to complementation of YFP fluorescence which is detectable in FACS-based assay format (Pan *et al*., 2021). We observed here low levels of YFP fluorescence when the CASK WT, E115K or N299S constructs were expressed alone in 293T cells. Coexpression of Liprin-α2 led to a strong increase in fluorescence for CASK-WT and the N299S variant, but not for the E115K variant (Fig. 10). These data suggest the existence of two different states for Liprin-α2: the Liprin-CASK-WT complex which is characterized by diffusely localized oligomers; and Liprin-α2 alone which forms LLPS-like droplets. The E115K variant fails to dissolve LLPS-based droplets into the more diffusely localized CASK-Liprin oligomers.

**Fig. 10.**
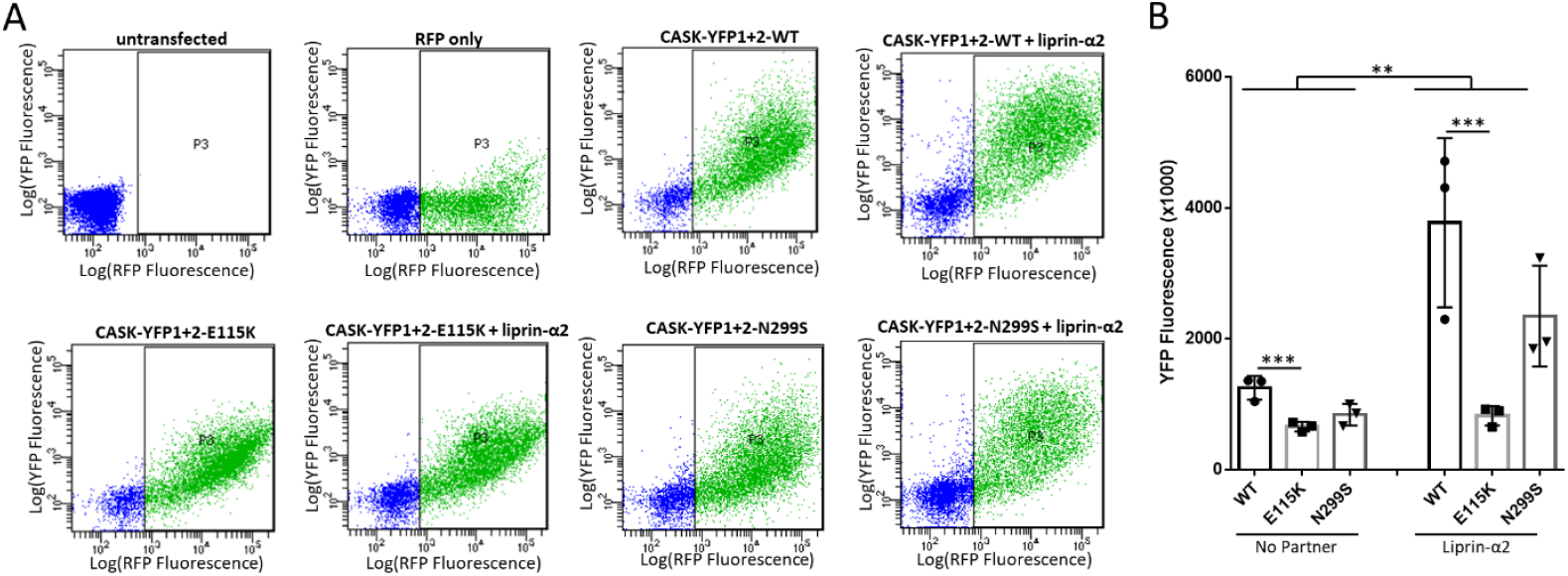
Liprin-α2 induces formation of CASK oligomers. **A**. HEK293T cells were transfected with plasmids coding for mRFP, CASK (WT or mutant) fused to the N- (YFP1) as well as the C-terminal (YFP2) halves of YFP, and Liprin-α2 as indicated. Two days after transfection, cells were harvested, resuspended in PBS and analyzed by flow cytometry using filters for mRFP and YFP fluorescence. **B**. Quantification of the data shown in A. Significance was determined by two-way ANOVA with Sidak’s multiple comparison test; **, p≤0.01; ***, p≤0.001; n=3. Mean ± *SD* is shown with each data point representing an independent transfection and flow cytometry experiment.

We sought additional proof that the dissolution of Liprin-α2 condensates by CASK is due to the direct interaction of CASK with Liprin. We made use of a mutation in the linker region between Liprin-α2 SAM domains 1 and 2, W981A. This alteration specifically disrupts binding to CASK, without disrupting other functions of the Liprin-α2 SAM domains (Wei *et al*., 2011). By coexpression of WT and mutant Liprin-α2 with CASK, followed by coimmunoprecipitation we confirmed that this substitution indeed eliminated the CASK-Liprin interaction (Fig. 11, A and B). In 293T cells, we observed that W981A mutant Liprin-α2 formed LLPS-like condensates very similar to the WT protein, and that coexpressed CASK-WT was unable to interfere with this cluster formation. Despite the complete loss of interaction seen in the biochemical experiment, CASK was recruited to these clusters where it extensively colocalized with Liprin-W891A (Fig. 11, C-E). We reproduced these results in cultured neurons, where the Liprin-α2 mutant formed LLPS-type condensates throughout the cell. Coexpression of CASK did not alter this localization, and CASK was again found in the same droplets as Liprin-α2 (Fig. 11, F-H). Thus, we conclude that, irrespective of the cell type, Liprin-α2 forms large condensates, and a tight interaction with CASK is required to regulate droplet formation.

**Fig. 11.**
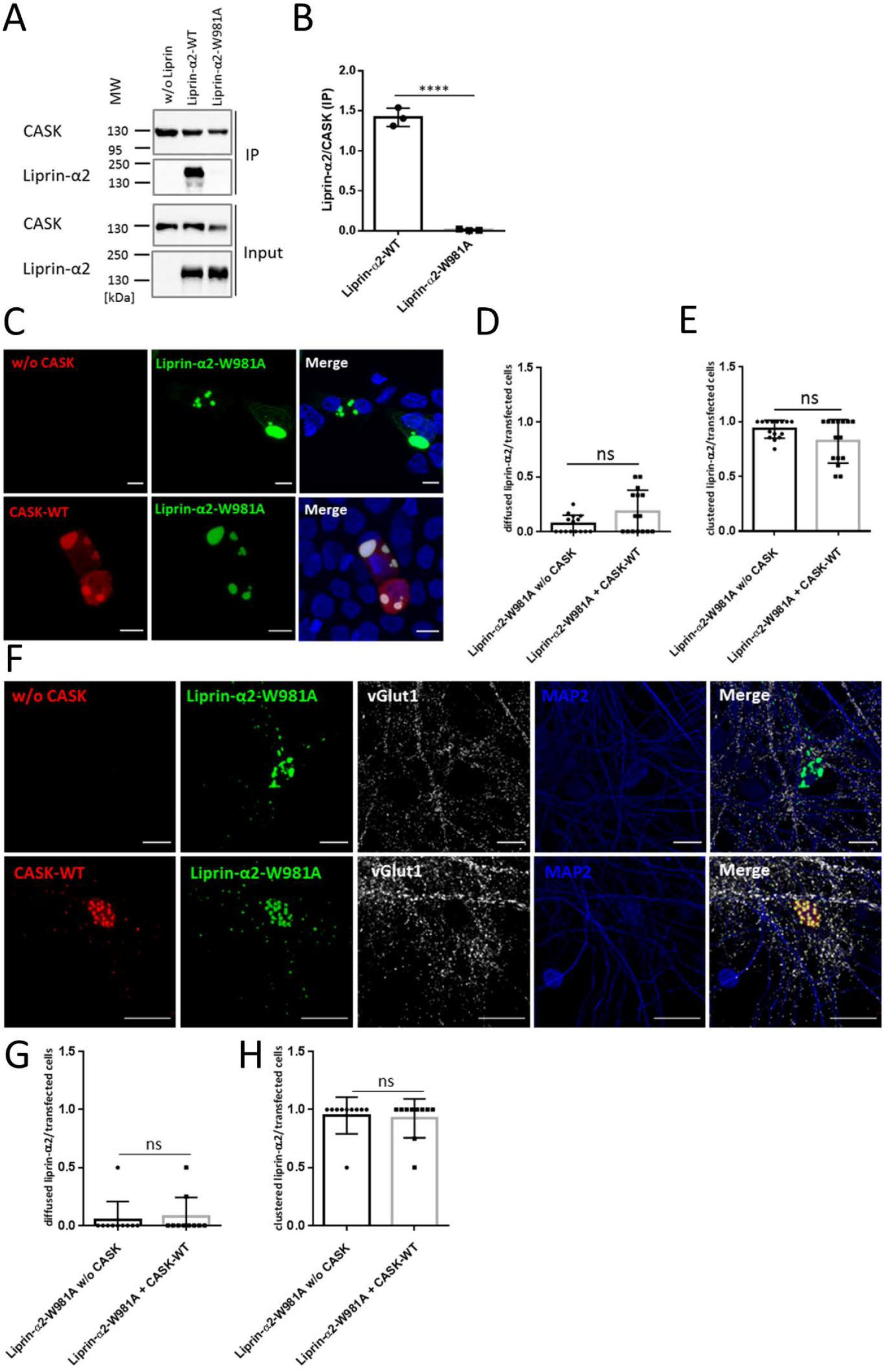
CASK needs to bind Liprin-α2 to negatively regulate condensate formation. **A**. mRFP-CASK was coexpressed with Liprin-α2-WT or the W981A mutant. After cell lysis and immunoprecipitation of CASK, input (IN) and precipitate (IP) samples were analysed by western blotting. **B**. Quantification of the data in shown in A with mean ±*SD* of three independent transfections depicted by single data points; n=3. **C**. Coexpression in HEK293T cells shows that W981A-mutant Liprin-α2 localizes to large intracellular clusters in the absence as well as in the presence of CASK-WT. **D, E**. Quantification of the cell populations shown in C based on the total number of transfected cells from 15 images. Shown is the mean ± *SD* with one data point per analysed image. **F**. Localization of W981A mutant Liprin-α2 and CASK-WT was analysed in primary hippocampal neurons, as before in Fig. 6. Here, CASK-WT did not alter the localizaton of the Liprin-α2 mutant. **G, H**. Quantification of hippocampal neurons as shown in F. 10 images per condition were analysed with up to five transfected neurons present and the mean ± *SD* is shown with each data point representing one analysed picture. Statistics were done with two-tailed Student’s *t*-test; ****, p≤0.0001; n=3-6.

## Discussion

We identified four male patients carrying *CASK* missense variants affecting the CaMK domain. All patients had a severe neurodevelopmental disorder, characterized by microcephaly, intellectual disability, and seizures. Patient 1, carrying the E115K variant, in addition showed pontine and cerebellar hypoplasia, a hallmark of the MICPCH phenotype described for *CASK* loss-of-function mutations (Moog *et al*, 2015; Najm *et al*., 2008). This patient died at a young age. By performing a thorough functional analysis of all four mutants, our goal was to determine which of the various functional aspects of the CASK CaMK domain is responsible for the patients’ phenotype. In addition, we wanted to find out what was special about the E115K variant, as it causes the additional severe PCH phenotype.

None of the variants appeared to affect folding or stability of overexpressed CASK protein. All four variants did not significantly affect binding to Mint1, to Veli proteins, and to Neurexin. Furthermore, none of the variants interfered with folding of the isolated CaMK domain prepared from bacteria. This allowed us to measure Mg^2+^-sensitive ATP binding. Mg^2+^-sensitivity instead of Mg^2+^-dependence classifies CASK as an atypical kinase (Mukherjee *et al*., 2008). Efficient binding was detected for all four mutants, similar to the wildtype, though variants R264K and N299S slightly altered the Mg^2+^ sensitivity of ATP binding. N299 is located in the kinase regulatory segment in the αR1 helix. A movement of this helix with respect to the rest of the domain, induced by the N299S variant, would change the geometry of the kinase active site, leading to altered binding parameters.

All four variants shared a strongly reduced ability to interact with Liprin-α2, suggesting that a weakened CASK-Liprin connection is responsible for the phenotype of all four patients. Together with our previous findings, showing that loss of Neurexin binding or Neurexin-induced oligomerization is a frequent result of pathogenic CASK missense variants (Pan *et al*., 2021), this points strongly to a presynaptic origin of CASK-related neurodevelopmental disorders. But what makes the E115K variant so devastating?

Liprin-α proteins have been shown to be early organizers of presynaptic development by recruiting ELKS, RIM and CASK (Dai *et al*, 2006; Spangler *et al*, 2013). The ability of Liprin-α proteins to undergo LLPS has now been documented in several studies; LLPS is driven by multimerization of its N-terminal coiled-coil motifs, which leads to a high local concentration of Liprin-α and associated ELKS molecules (Liang *et al*., 2021). Furthermore, a central intrinsically disordered region (see Fig. 5, C) in Liprin-α proteins from different species contributes to LLPS (Emperador-Melero *et al*., 2021; McDonald *et al*., 2020). We observed here that CASK has a regulatory effect on condensate formation. This depends on the direct interaction of the two proteins, as it can be abolished by mutations E115K in CASK and W981A in Liprin-α2. Structurally, we do not know how interaction with CASK negatively affects condensation of Liprin-α2 into droplets. CASK binds to the C-terminal part of Liprin-α2 which has so far not been implicated in condensation. One aspect may be phosphorylation of the N-terminal coiled-coil domain of Liprin-α2 at Ser87, which is reduced upon CASK binding. LLPS of Liprin-α3 is triggered by a phosphorylation event, in this case within the IDR region at Ser760, through the activity of protein kinase C (Emperador-Melero *et al*., 2021). As S760 is absent in Liprin-α2, and S87 is absent in Liprin-α3, condensation appears to be differentially regulated by phosphorylation events.

Upon CASK binding, a long α-helical segment (αN-segment in Figs. 1, 5) immediately N-terminal to the SAM domains of Liprin-α performs a significant outward turn (Wei *et al*., 2011; Xie *et al*., 2021). As the αN-segment is close in sequence to the intrinsically disordered region (IDR), it is conceivable that this movement of αN alters the propensity of the IDR to condensate and induce phase separation.

Data from *C. elegans* suggest that LLPS mediated by Liprin is an essential step in synapse formation; however, the Liprin aggregates formed during LLPS are modified at a later stage in synaptogenesis, leading to some form of solidification of the active zone (McDonald *et al*., 2020). In this respect, our findings that CASK can dissolve preformed LLPS like clusters of Liprin-α2 points to a mechanism where LLPS is required for early stages of active zone formation, whereas during a later stage CASK and possibly Neurexin are added to the complex. This likely occurs in the form of CASK oligomers, as depicted by our split-YFP data that would then allow for restructuring of the large Liprin-based condensates.

Importantly, the inability to regulate phase transitions of Liprin-α2 is the single functional feature which distinguishes the CASK E115K variant from the other three investigated variants. E115K caused a PCH phenotype with early lethality, whereas the other three variants did cause a severe neurodevelopmental disorder but without PCH. In further studies it will be important to delineate how the aberrant regulation of LLPS-like condensate formation by Liprin-α proteins contributes to pontocerebellar hypoplasia.

## Materials and Methods

### Patients and genetic analysis

We identified four patients with a *CASK* missense variant from different diagnostic and research cohorts from across India and the United States. Genetic testing was performed by Sanger sequencing of *CASK*, targeted next-generation sequencing gene panel or exome sequencing. Clinical and molecular findings in patients 1 to 4 are summarized in Table 1. Informed consent for genetic analysis was obtained from parents/legal guardians, and genetic studies were performed clinically or as approved by the Institutional Review Boards of the respective institution.

Patients for this study were ascertained over a course of two years. Therefore, in some panels of our functional assays, only one or two variants are compared with the respective wild type condition.

### Expression constructs

For expression in HEK293T cells, cDNA coding for CASK transcript variant 3 (TV3; (Tibbe *et al*, 2021)) fused to an N-terminal mRFP-tag in pmRFP-N1 was used. For expression in neurons, cDNAs coding for mRFP-CASK-TV5 fusion proteins were inserted into a vector carrying the human synapsin promoter (Repetto *et al*, 2018; Tibbe *et al*., 2021). For the preparation of fusion proteins, the cDNA encoding the CaMK domain was cloned into the pET-SUMO vector coding for a His_6_-SUMO tag (Thermo Scientific). An expression vector for GFP-Mint1 was obtained from C. Reissner and M. Missler (Münster, Germany). HA-tagged Neurexin-1β was from P. Scheiffele (Basel, Switzerland) via Addgene (58267), HA-tagged and GFP-tagged Liprin-α2 were from C. Hoogenraad (Utrecht, The Netherlands). Mutations were introduced using the Quik-Change II site-directed mutagenesis kit (Agilent), using two complementary, mutagenic oligonucleotides. Constructs were verified by Sanger sequencing.

### Antibodies

For western blotting experiments, all primary antibodies were diluted 1:1,000 in TBS-T with 5% milk powder (MP). The following antibodies were used: α-CASK (Rb, Cell Signaling Technologies, #9497S), α-GFP (Ms, Covance, #MMS-118P), α-Veli 1/2/3 (Rb, Synaptic Systems, #184 002), α-HA (Ms, Sigma, #H9658) and α-Myc (Ms, Sigma, #M5546). Horse radish peroxidase (HRP)-coupled secondary antibodies were used in a dilution of 1:2,500 in TBS-T (Gt-α-Ms or Gt-α-Rb, ImmunoReagents, #BOT-20400 or #BOT-20402). For application in immunocytochemistry of hippocampal neurons, two primary antibodies, both prepared in 1:1000 dilutions in 2% horse serum (HS) in PBS, were used: α-MAP2 (Ck; Antibodies Online, #ABIN 111 291) and α-vGlut1 (Rb; Synaptic Systems #135 303). As secondary antibodies we used Alexa-405 (Gt-α-Ck, Abcam #ab175675) and Alexa-633 (Gt-α-Rb, Thermo Fisher, #A-21071).

### Cell culture, transfection and coimmunoprecipitation

HEK293T cells (ATCC®, CRL-3216™) were cultivated on 10 cm dishes in Dulbecco’s Modified Eagle Medium (DMEM) supplemented with 1 x penicillin/streptomycin and 10% fetal bovine serum at 37 °C, 5% CO_2_ and humidified air. Cells were transiently transfected with TurboFect transfection reagent (Thermo Fisher) according to the manufacturer’s instructions. 24 h after transfection, the cells were washed in PBS and lysed in 1 ml of RIPA buffer supplemented with protease inhibitors (0.125 M phenylmethylsulphonyl fluoride, 5 mg/mL leupeptin, 1 mg/mL pepstatin A). Lysates cleared by centrifugation (15 min, 4 °C at 20,000 × g) were subjected to immunoprecipitation with 20 µL of RFP-Trap agarose beads (ChromoTek, Munich, Germany) for 2 h at 4 °C under rotation. The beads were washed five times with RIPA buffer, followed by centrifugation (1 min, 4 °C, 1000 × g) and immunoprecipitate (IP) and input samples (IN) were processed for western blotting. Protein bands were detected by chemoluminescence with a BioRad imaging system in the “auto-mode”, avoiding over-saturation while maximizing signal intensity. Band intensities were quantified using ImageLab 6.0 software.

### Bacterial expression and purification of fusion proteins

His_6_-SUMO-tagged fusion proteins were expressed in BL21 (DE3) cells and purified from bacterial lysates prepared in native lysis buffer (50 mM NaH_2_PO_4_, 500 mM NaCl, pH 8.0) using Ni–NTA agarose (Qiagen, Hilden, Germany). Proteins were eluted from beads with 250 mM imidazole in lysis buffer and were immediately applied to G-25 columns equilibrated in TNP-ATP-binding buffer (40 mM Tris HCl 100 mM NaCl, 50 mM KCl, pH 7.5), followed by elution in the same buffer. Efficiency of protein purifications was verified by SDS-PAGE, followed by Coomassie staining. Protein concentrations were determined by Bradford assay, using BSA as a standard.

### TNP-ATP binding assay

The binding of 2,4,6-trinitrophenol conjugated ATP (TNP-ATP) to the CaMK domain of CASK was measured by fluorescence spectroscopy. 1 μM TNP-ATP was added to 100 µg/mL soluble protein in TNP-ATP binding buffer. For the detection of magnesium sensitivity, 2 mM Mg^2+^ was added. After 15 min incubation, the fluorescence emission spectra from 500 nm to 600 nm was measured on a Synergy H1 plate reader.

### Cell culture of primary hippocampal neurons

Primary cultures of hippocampal neurons were prepared from *Rattus norvegicus* embryonic day 18 (E18) rats (Wistar Unilever outbred rat, strain: HsdCpb:WU; Envigo) regardless of gender, as described before (Hassani Nia *et al*, 2020). Neurons were isolated using papain neuron isolation enzyme (Thermo scientific, #88285) and cultivated in neurobasal culture media containing B27 and GlutaMAX supplements (Thermo scientific, #21103049, #17504044 and #A1286001). Neurons were transfected using the calcium phosphate method after 7 days in vitro (DIV7), as described before (Hassani Nia *et al*., 2020). Cells were cultured until DIV14 before fixation and staining.

Animal experiments were approved by, and conducted in accordance with, the guidelines of the Animal Welfare Committee of the University Medical Center (Hamburg, Germany) under permission number Org1018.

### Immunocytochemistry and confocal microscopy

Transfected HEK293T cells were transferred to PLL-coated coverslips in 12 well plates one day before fixation. Hippocampal neurons were cultured and transfected on PLL-coated coverslips in 12 well plates. For ICC, cells were washed three times with PBS and fixed in 4% PFA with 4% Sucrose in PBS for 15 min at room temperature (RT). The cells were washed again three times with cold PBS, followed by permeabilization in 0.1% Triton-X 100 in PBS for 3 min at RT. After washing with PBS, cells were incubated in 10% HS in PBS for 1 h at RT to reduce unspecific antibody binding. Primary antibodies were prepared in 2% HS in PBS and the cells were incubated with the antibody solution overnight at 4 °C in a humidified atmosphere. After washing with PBS, cells were incubated with secondary antibodies diluted in PBS for 1 h at RT. After washing three times with PBS and once with ddH2O, coverslips were mounted with ProLong Diamond Antifade mounting medium (Thermo Fisher, #P36961). Samples were analysed by confocal microscopy, using a Leica SP8 confocal microscope (provided by UKE Microscopy Imaging Facility; UMIF).

### FastAP dephosphorylation assay

HEK293T cells expressing HA-Liprin-α2 alone or together with mRFP-CASK-wildtype (WT) were lysed in IP buffer without EDTA. Cleared lysates were treated with or without FastAP Thermosensitive Alkaline Phosphatase (Thermo Scientific, EF0651), as described in the manufacturer’s protocol for protein dephosphorylation. After incubation at 37 °C for 1 h under shaking, samples were analysed by immunoblotting.

### Sample preparation for proteome analysis

HA-tagged Liprin-α2 was immunoprecipitated from transfected cells using HA-specific magnetic beads. After washing, samples were diluted in 1% w/v sodium deoxycholate (SDC) in 100 mM triethyl bicarbonate buffer and boiled at 95 °C for 5 min. Disulfide bonds were reduced in the presence of 10 mM dithiothreitol (DTT) at 60 °C for 30 min. Cysteine residues were alkylated in presence of 20 mM iodoacetamide at 37 °C in the dark for 30 min and tryptic digestion (sequencing grade, Promega) was performed at a 100:1 protein to enzyme ration at 37 °C over night. Digestion was stopped and SDC precipitated by the addition of 1% v/v formic acid (FA). Samples were centrifuged at 16,000 g for 5 min and the supernatant was transferred into a new tube. Samples were dried in a vacuum centrifuge.

### LC-MS/MS in Data Dependent mode

Samples were resuspended in 0.1% formic acid (FA) and transferred into a full recovery autosampler vial (Waters). Chromatographic separation was achieved on a UPLC system (nanoAcquity, Waters) with a two-buffer system (buffer A: 0.1% FA in water, buffer B: 0.1% FA in ACN). Attached to the UPLC was a C18 trap column (Symmetry C18 Trap Column, 100Å, 5 µm, 180 µm x 20 mm, Waters) for online desalting and sample purification followed by an C18 separation column (BEH130 C18 column, 75 µm x 25 cm, 130 Å pore size, 1.7 µm particle size, Waters). Peptides were separation using a 60 min gradient with increasing acetonitrile concentration from 2% -30%. The eluting peptides were analyzed on a quadrupole orbitrap mass spectrometer (QExactive, Thermo Fisher Scientific) in data dependent acquisition (DDA).

### Data analysis and processing

Acquired DDA LC-MS/MS data were searched against the reviewed human protein database downloaded from Uniprot (release April 2020, 20,365 protein entries, EMBL) using the Sequest algorithm integrated in the Proteome Discoverer software version 2.4 (Thermo Fisher Scientific) in label free quantification mode with match between runs enabled, performing chromatographic retention re-calibration for precursors with a 5 min retention time tolerance, no scaling, and no normalization for extracted peptide areas was done. Mass tolerances for precursors was set to 10 ppm and 0.02 Da for fragments. Carbamidomethylation was set as a fixed modification for cysteine residues and the oxidation of methionine, phosphorylation of serine and threonine, pyro-glutamate formation at glutamine residues at the peptide N-terminus as well as acetylation of the protein N-terminus, methionine loss at the protein N-terminus and the acetylation after methionine loss at the protein N-terminus were allowed as variable modifications. Only peptide with a high confidence (false discovery rate < 1% using a decoy data base approach) were accepted as identified.

### Split-YFP experiments

HEK293T cells were cotransfected with 3 μg of CASK-YFP1 and CASK-YFP2 expression vectors, in combination with 1 μg of pmRFP-C1 plasmid and either 3 μg of Liprin-α2 or empty vector. Two days later, cells were trypsinized, centrifuged at 1,000 x g for 5 min, and resuspended in PBS. Flow cytometry was performed at a FACS Canto-II instrument (BD Biosciences, Heidelberg, Germany). Transfected cells were identified by mRFP fluorescence, and YFP fluorescence was quantified from 10,000 fluorescence events from viable single cells for each condition. Aliquots of cells were additionally analysed by immunoblotting to determine efficient expression of CASK fusion proteins.

## Acknowledgements

We thank Hans-Hinrich Hönck for excellent technical support, UKE microscoping imaging facility (umif) for providing microscopes, Carsten Reissner and Markus Missler (Münster, Germany), Peter Scheiffele (Basel, Switzerland) and Caspar Hoogenraad (Utrecht, The Netherlands) for plasmids. We thank the UKE microscopy imaging facility (umif) for providing confocal microscopes. This work was supported by grants from Deutsche Forschungsgemeinschaft (KU 1240/10-1 to K.K., KR 1321/7-1 to H.-J.K.), and by Jordan’s Guardian Angels, the Sunderland Foundation and the Brotman Baty Institute (to G.M.M.).

